# L-2-hydroxyglutarate recycling is linked to coenzyme Q biosynthesis

**DOI:** 10.64898/2026.07.16.738879

**Authors:** Marco Malatesta, Andrea Gottinger, Callum R. Nicoll, Diran Herebian, Demetra Mattiroli, Domiziana Cecchini, Annette Seibt, Andrea Alfieri, Felix Distelmaier, Andrea Mattevi

**Affiliations:** Department of Biology and Biotechnology ‘Lazzaro Spallanzani’, University of Pavia, Via Ferrata 9, 27100, Pavia, Italy; Department of General Pediatrics and Neonatology, Medical Faculty and University Hospital Düsseldorf, Heinrich-Heine-University, Düsseldorf, Germany; Centro Grandi Strumenti, University of Pavia, Pavia, Italy

## Abstract

The mitochondrial COQ metabolon catalyzes the late stages of the biosynthesis of coenzyme Q, an essential and ubiquitous cofactor. Here, by integrating coevolution, coexpression, colocalization and domain-fusion analyses, we identify L-2-hydroxyglutarate dehydrogenase (L2HGDH) as an integral component of this assembly. By acting in physical proximity to the COQ metabolon, L2HGDH sustains coenzyme Q production by maintaining the biosynthetic intermediates in their catalytically-active reduced state. Consistently, analysis of fibroblasts and urine samples from patients with primary coenzyme Q deficiency displayed marked accumulation of L-2-hydroxyglutarate. Cryo-electron microscopy reveals that L2HGDH forms a stable complex with COQ3 and COQ6, defining a heterotrimeric assembly that organizes catalytic sites on a shared membrane-facing surface thereby enabling localized quinone reduction. Together, these findings identify L2HGDH as a previously unrecognized component of the COQ metabolon, establish a direct link between central carbon metabolism and coenzyme Q biosynthesis, and expand the functional roles of metabolons in coordinating metabolic flux across distinct pathways.

## Introduction

Metabolon, a term coined more than four decades ago, describes supramolecular assemblies of enzymes that enhance pathway efficiency through dynamic physical interactions^1,2^. This concept is illustrated by several widely studied systems. Two paradigmatic cases are the tricarboxylic acid (TCA) cycle^3,4^ and the *purinosome*^5,6^, in which enzymes are spatially coordinated through transient interactions to facilitate substrate channeling. Beyond improving catalytic efficiency, such spatial arrangements reflect the extensive interconnection of cellular metabolism. The TCA cycle, for instance, serves as a central hub connecting multiple metabolic pathways through shared enzymes and branching points^7^: succinate dehydrogenase is also part of the electron transport chain^8^, whereas malate dehydrogenase operates in the malate-aspartate shuttle that mediates the interconversion of malate and oxaloacetate across mitochondrial and cytosolic compartments^9^.

Another example is represented by the so-called *COQ metabolon* that comprises a core of six proteins (COQ3, COQ4, COQ5, COQ6, COQ7, and COQ9) located on the matrix side of the inner mitochondrial membrane^10^ (**Figure 1A**). Although coenzyme Q biosynthesis has been studied for decades^11–13^, recent structural and biochemical studies have progressively revealed that this assembly does not organize in a closed module, but rather in a dynamic assembly that interfaces with multiple mitochondrial processes through auxiliary factors. Specifically, the atypical kinase COQ8 facilitates intermediate trafficking along the pathway^14,15^; FDXR and FDX2 connect coenzyme Q biosynthesis to the mitochondrial ferredoxin system and iron-sulfur cluster biogenesis by providing reducing equivalents to COQ6^10,16^; PYURF stabilizes COQ5 and enhances its enzymatic function while also supporting NDUFAF5, an assembly factor for respiratory complex I^17^; and RTN4IP1 maintains the reduced state of quinone intermediates required for methyltransferase activity^18^. These observations raise the question of whether additional, as-yet-unidentified components contribute to the architecture of this assembly and mediate crosstalk with other metabolic pathways.

**Figure 1.**
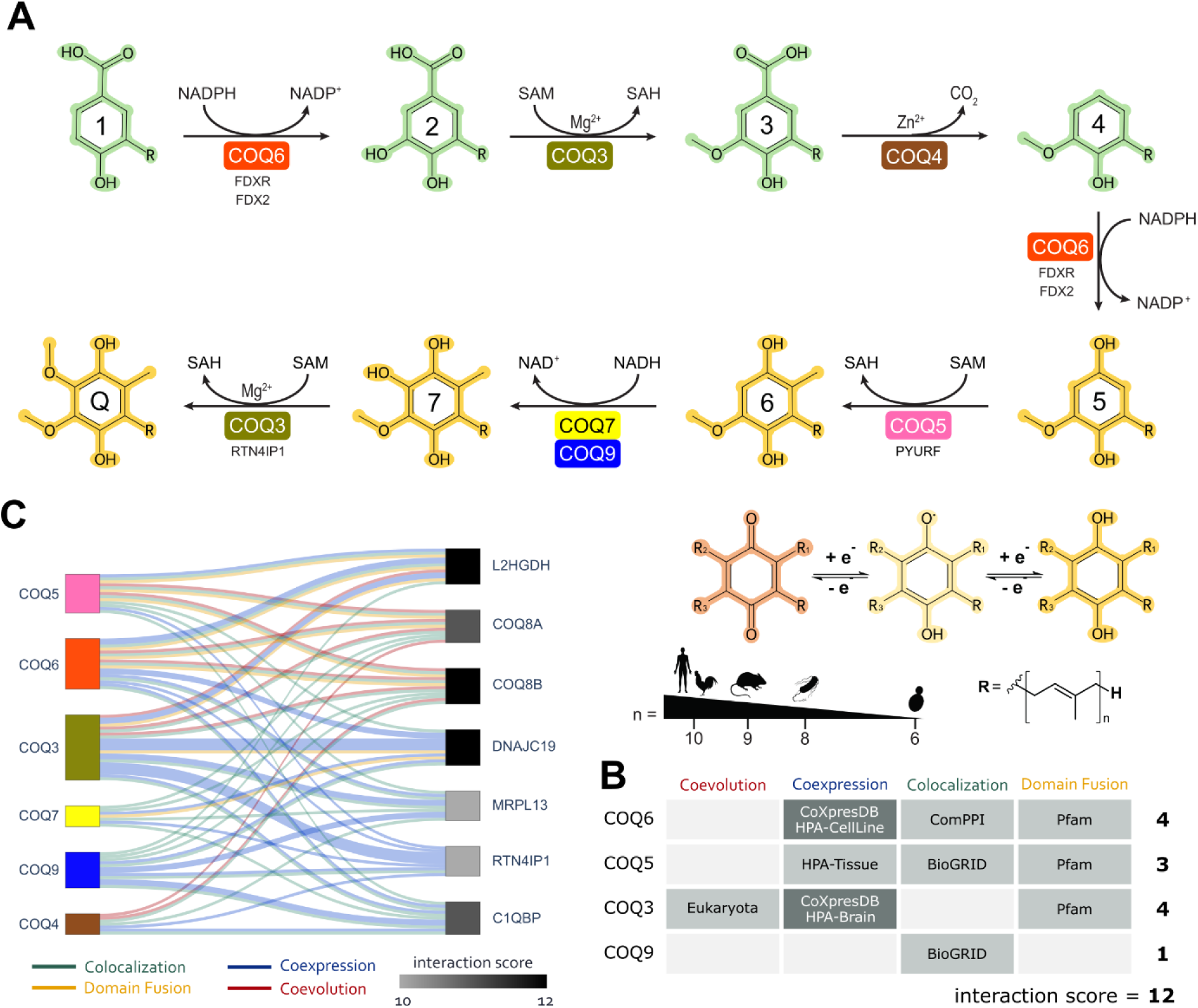
Identification of L2HGDH as a candidate partner of the COQ metabolon (A) Schematic representation of coenzyme Q biosynthesis. Intermediates are numbered from **1**, prenylated *p*-hydroxybenzoate, to **7** whereas the final product (coenzyme Q) is indicated as **Q**. Core metabolon components are shown in distinct colors, while accessory proteins are indicated in smaller text below their corresponding enzymatic step. Non-quinone intermediates are highlighted in green, whereas reduced quinone intermediates of the second half of the pathway are highlighted in yellow. These latter intermediates can undergo redox reactions, as schematized at the bottom. Organism-specific variations in polyisoprenoid tail length are also indicated. (B) Functional association summary between L2HGDH and COQ metabolon components. Cell intensity reflects the number of positive hits for each evidence source. (C) Sankey plot depicting the interactions within the human COQ metabolon. Core components (COQ3-COQ7:9) were used as seed nodes to integrate coevolution, coexpression, colocalization, and domain-fusion. Links represent interactions between COQ proteins and candidate partners, with widths proportional to the interaction score shown as a grayscale gradient.

Here, we systematically explored candidate interactors of the human COQ metabolon by integrating multiple layers of functional association data. This approach uncovered previously uncharacterized candidates from which emerged L2HGDH, a mitochondrial FAD-dependent dehydrogenase involved in the recycling of the oncometabolite L-2-hydroxyglutarate^19,20^. Through a combination of biochemical reconstitution and cryo-electron microscopy, we demonstrate that L2HGDH acts as a quinone-dependent reductase that functionally and physically integrates into the COQ metabolon while simultaneously recycling a metabolic byproduct leaked from the TCA cycle. Measurements of hydroxyglutarate levels in COQ4-deficient patient samples display altered L-2-hydroxyglutarate homeostasis, providing evidence for a physiological link between coenzyme Q biosynthesis and L-2-hydroxyglutarate metabolism. Our findings reveal an integrated and organized redox network linking coenzyme Q biosynthesis, central carbon metabolism, and L-2-hydroxyglutarate recycling.

## Results

### Functional interaction network identifies L2HGDH as candidate partner of the COQ metabolon

To identify novel protein associations with the human COQ metabolon, we first built a functional interaction network across the human proteome by integrating data from four curated and validated databases: coevolution^21,22^, coexpression (Human Protein Atlas^23^ and COXPRESdb^24^), colocalization (ComPPI^25^), and domain fusion (Pfam^26^). These databases were queried to identify possible structural and functional interactions between COQs and all proteins of the human proteome. As detailed in the **Methods** section, the putative interactors of the COQ metabolon were ranked based on the total number of hits (see **Figure 1B** for an example). Proteins with an interaction score above ten included several established regulators of the metabolon (**Figure 1C**). Notably, COQ8A, COQ8B and RTN4IP1 are known for their roles in metabolon activity and coenzyme Q biosynthesis^10,18^. Several additional high-scoring candidates, however, lacked any previous direct association with coenzyme Q biosynthesis. Among these, DNAJC19 emerged as one of the strongest nodes in the network. DNAJC19 is a mitochondrial co-chaperone involved in cardiolipin remodeling through interactions with prohibitins^27^, consistent with the strong functional relationship between cardiolipin and the COQ metabolon, where this lipid stimulates COQ8 activity^14,15^. Additional candidates included C1QBP, a multifunctional mitochondrial matrix protein involved in mitochondrial translation^28^, and MRPL13, whose *Saccharomyces cerevisiae* ortholog is associated with membrane-bound mitoribosomes responsible for the translation of mitochondrial proteins^29^. In most cases, these associations were mainly supported by coexpression and colocalization.

Notably, L-2-hydroxyglutarate dehydrogenase (L2HGDH) was the only candidate supported by all four evidence sources (**Figures 1B-C**). L2HGDH is a mitochondrial FAD-dependent oxidoreductase that catalyzes the oxidation of L-2-hydroxyglutarate to α-ketoglutarate^19,20^. L-2-hydroxyglutarate is a metabolic byproduct that accumulates under hypoxic conditions due to the promiscuous activity of lactate dehydrogenase and malate dehydrogenase on α-ketoglutarate^30,31^. Of note, α-ketoglutarate is a TCA-cycle intermediate and obligatory cofactor of a number of epigenetic and regulatory enzymes^32–34^. Loss-of-function mutations in L2HGDH cause L-2-hydroxyglutaric aciduria, a rare neurometabolic disorder characterized by the accumulation of L-2-hydroxyglutarate. To validate our prediction, we explored a recent crosslinking mass spectrometry dataset of the mitochondrial membrane spatial proteome^35^. This analysis recovered crosslinks between COQ3 and COQ6, the enzymes catalyzing the first two steps of head-group modification during coenzyme Q biosynthesis (**Figure 1A**) and revealed additional crosslinks connecting L2HGDH with both COQ3 and COQ6. We concluded that L2HGDH might represent *a bona fide* structural and functional partner of the COQ metabolon.

### L2HGDH utilizes coenzyme Q biosynthetic intermediates as electron acceptors

To experimentally investigate the potential roles of L2HGDH in coenzyme Q biosynthesis, we reconstructed an ancestral tetrapod variant of the enzyme using ancestral sequence reconstruction, similarly to what was previously established for other COQ proteins^10^. This strategy ensures evolutionary consistency across the reconstructed system and preserves potential interaction interfaces. The reconstructed tetrapod L2HGDH shares 80% sequence identity with the human enzyme, substantially higher than the 56% sequence identity of the previously structurally characterized *Drosophila* ortholog^36^ (**Figure S1A**). Tetrapod L2HGDH was expressed in *E. coli* and purified in sufficient yield for structural and biochemical studies (**Figures S1B-C**). The recombinant enzyme fully retained its FAD cofactor suggesting proper folding and functional competence (**Figure 2A**). We therefore performed various experiments to investigate its biochemical and enzymatic properties.

**Figure 2.**
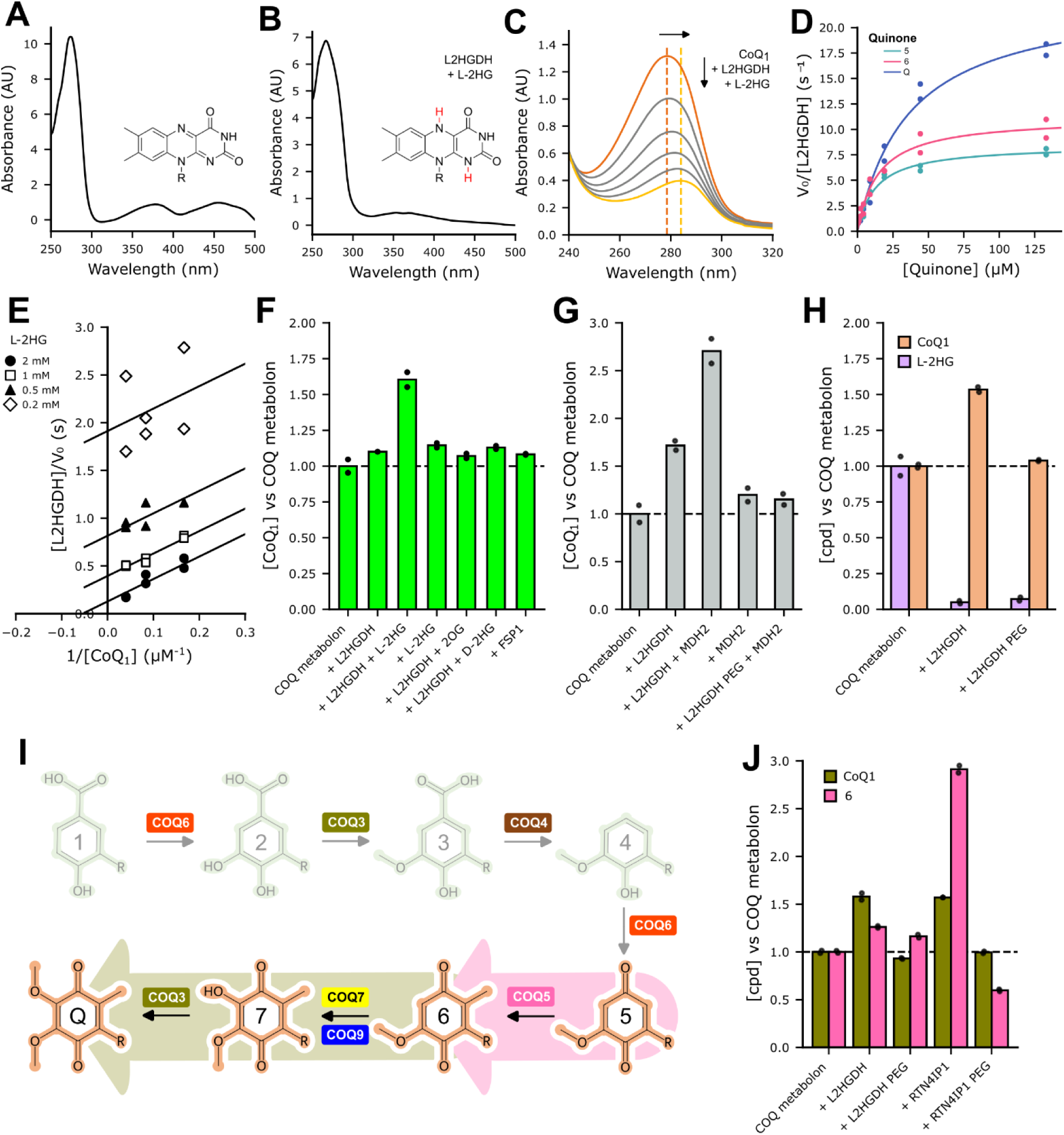
L2HGDH functions as a quinone reductase that promotes coenzyme Q biosynthesis. (A) UV-visible absorption spectrum of recombinant L2HGDH (250 μM), showing the characteristic oxidized FAD signal with its typical double peak. (B) UV-visible absorption spectrum of L2HGDH in the presence of L-2-hydroxyglutarate (L-2HG; 2 mM). The FAD peak disappears due to the lack of reoxidation by molecular oxygen, confirming that L2HGDH does not function as an oxidase. (C) UV-visible spectra of 100 μM coenzyme Q_1_ in reaction buffer (20 mM HEPES pH 7.5, 50 mM NaCl). Upon addition of 100 nM L2HGDH and 2 mM L-2HG, a decrease in absorbance and a spectral shift toward 285 nm are observed, consistent with quinone reduction. Dashed lines indicate the absorption maxima of the oxidized (salmon) and reduced (yellow) forms. Spectra were collected every 30 s as a time course of the reaction. (D) Nonlinear Michaelis-Menten fits with quinone substrates from the coenzyme Q biosynthetic pathway (**5**, **6** and coenzyme Q_1_). Each data point represents measurements from a single independent experiment (n=2). Reactions were performed in 25 mM HEPES (pH 7.4) in the presence of L-2-hydroxyglutarate, the indicated quinone substrate, glutamate dehydrogenase, NADH, NH_4_Cl and L2HGDH. (E) Double-reciprocal (Lineweaver-Burk) plots obtained by varying coenzyme Q concentrations at fixed concentrations of L-2-hydroxyglutarate (0.2, 0.5, 1, and 2 mM). Symbols indicate individual measurements, whereas lines correspond to global linear fits. The approximately parallel lines are indicative of a ping-pong kinetic mechanism involving sequential binding and dissociation of L-2-hydroxyglutarate and quinone substrate, respectively. (F) Bar plot showing the fold change in coenzyme Q_1_ production by the complete metabolon (COQ3, COQ4, COQ5, COQ6, COQ7, COQ8B, COQ9, FDX2, FDXR, their cofactors/co-substrates NADP^+^, NAD^+^, SAM, ATP, Zn^2+^, the precursor **1**, and glucose dehydrogenase with glucose for NAD(P)H regeneration) upon addition of the components indicated on the x-axis. Quantification was performed by LC-MS using integrated peak areas normalized to the internal standard (sorbicillin). (G) Bar plot showing the fold change in coenzyme Q_1_ production. The complete metabolon is used as reference control (COQ metabolon), and addition of pig-heart malate dehydrogenase (MDH2) and L2HGDH are indicated on the x-axis. (H) Bar plot showing the fold change in coenzyme Q_1_ production (salmon) and L-2-hydroxyglutarate consumption (lilac) upon addition of L2HGDH or its PEGylated variant (L2HGDH PEG). (I) Schematic representation of the coenzyme Q biosynthetic pathway and the enzymes involved according to the KEGG database. Intermediates are numbered from **1** to **7**, and the final product coenzyme Q is abbreviated as Q. Compounds are colored to distinguish the early non-quinone phase of the pathway (green) from the late quinone phase (salmon), beginning after the second reaction catalyzed by COQ6. Arrows are colored according to the reactions monitored in the experiments shown in panel **J**. (J) Bar plot showing the fold change in product formation for the reactions illustrated in panel **I**, in the presence of the complete metabolon upon addition of L2HGDH and RTN4IP1 in their native or PEGylated form as indicated on the x-axis. To monitor the COQ5-dependent reaction (pink), **5** was used as substrate and **6** was quantified as product. The reaction mixture was depleted of NAD^+^ to prevent consumption of **6** by COQ7. To monitor the COQ3-dependent reaction (olive), **6** was used as substrate and coenzyme Q_1_ was quantified as product.

To assess electron acceptor specificity, we tested NAD⁺, NADP⁺, and molecular oxygen as alternative oxidants. None of them supported catalytic turnover, indicating that L2HGDH functions neither as a classical NAD(P)H-dependent oxidoreductase nor as an oxidase (**Figure 2B**). In agreement with recent literature^19,37^, we then hypothesized that the enzyme could exploit oxidized quinones of the biosynthetic pathway as electron acceptors. Maintaining quinone biosynthetic intermediates in a reduced state could support the methyltransferase reactions of the pathway, similarly to what has been proposed for RTN4IP1^18^ (**Figure 1A**). We first monitored the reduction of the soluble mono-prenylated coenzyme Q (coenzyme Q_1_; CoQ_1_) spectrophotometrically. Upon addition of L2HGDH and L-2-hydroxyglutarate, we observed a spectral shift toward 290 nm, consistent with quinone reduction (**Figure 2C**). We next asked whether L2HGDH could also use quinone intermediates of the coenzyme Q biosynthetic pathway as electron acceptors. Following the second hydroxylation step, pathway intermediates enter a sort of “quinone phase” as they become capable of undergoing redox cycling. Indeed, the two methyltransferase reactions of the metabolon, COQ3 and COQ5, strictly depend on the availability of reduced substrates^38,39^ (**Figure 1A**). Because compound **7** could not be chemically synthesized, we restricted our screening to coenzyme Q_1_**, 5** and **6**. α-ketoglutarate formation was coupled to glutamate dehydrogenase activity and continuously monitored through NADH oxidation at 340 nm (**Figure S1D**). This assay confirmed efficient catalytic turnover with coenzyme Q_1_ and the other pathway intermediates (**Figure 2D**). No strong substrate preference was observed, as catalytic efficiencies (*k_cat_*/*K_M_*) were comparable among substrates, although *k_cat_* values were slightly lower for **5** and **6** compared to coenzyme Q_1_ besides their better *K_M_* values (**Figure 2D; Table S1**). To gain mechanistic insight into quinone reduction by L2HGDH, we performed steady-state kinetic analyses varying coenzyme Q concentrations at fixed levels of L-2-hydroxyglutarate. Double-reciprocal (Lineweaver-Burk) plots yielded a series of approximately parallel lines (**Figure 2E**). These data are consistent with a ping-pong mechanism, in which L-2-hydroxyglutarate first reduces the bound FAD cofactor and is released as α-ketoglutarate, followed by the binding of the quinone electron acceptor that re-oxidizes the flavin.

### L2HGDH enhances coenzyme Q biosynthesis through physical integration into the COQ metabolon

We next examined the impact of L2HGDH within a fully reconstituted coenzyme Q biosynthetic system. In reactions containing monoprenylated 4-hydroxybenzoate (**1**), the reconstructed COQ metabolon (COQ3-9), cofactors, and regeneration systems, addition of L2HGDH together with L-2-hydroxyglutarate resulted in an approximately 1.6-fold increase in coenzyme Q_1_ production (**Figure 2F**). This enhancement was strictly dependent on both enzyme and substrate, as neither L2HGDH nor L-2-hydroxyglutarate alone stimulated production. Because mitochondrial production of L-2-hydroxyglutarate predominantly originates from malate dehydrogenase reducing α-ketoglutarate, we next employed a regeneration system using pig-heart malate dehydrogenase. Under these conditions, coenzyme Q production increased up to ∼2.7-fold, showing that continuous regeneration of L-2-hydroxyglutarate can further stimulate coenzyme Q biosynthesis (**Figure 2G**). The effect was further confirmed by control experiments, including the addition of the reaction product α-ketoglutarate and the replacement of L-2-hydroxyglutarate with its D enantiomer, neither of which supported the observed stimulatory effect (**Figure 2F**). When an unrelated quinone reductase (FSP1) was substituted for L2HGDH, coenzyme Q levels remained comparable to the control, despite FSP1 displaying sustained reductase activity toward pathway intermediates in biochemical assays (**Figures 2F** and **S2; Table S1**). This observation suggested that quinone reductase activity alone is insufficient to support coenzyme Q biosynthesis and that an additional level of functional coupling is required. To test whether physical proximity and direct interaction with the metabolon could account for this effect, we chemically labeled the surface of L2HGDH using methoxy-polyethylene glycol (PEG), a treatment expected to sterically hinder protein-protein interactions as used previously for the same purpose^40^. Size-exclusion chromatography of PEGylated L2HGDH showed the expected shift toward larger apparent molecular species compared to the native protein (**Figure S3A**), while PEGylation did not affect its enzymatic activity toward quinone substrates (**Figure S3B**). However, PEG-L2HGDH completely lost the ability to stimulate coenzyme Q production within the metabolon, despite maintaining L-2-hydroxyglutarate consumption rates comparable to the native enzyme (**Figures 2G-H**). Together, these findings demonstrated that quinone reductase activity alone is insufficient and that physical integration within the metabolon is required to sustain local quinone reduction and efficient coenzyme Q production. Reduced quinone intermediates generated in solution are likely to undergo rapid spontaneous reoxidation unless they remain in close proximity to the enzyme that immediately consumes them during biosynthesis.

As a comparison, we investigated RTN4IP1, a quinone reductase recently demonstrated to support coenzyme Q biosynthesis by maintaining reduced quinone intermediates^18^. To evaluate its role within our reconstructed metabolon, we generated the ancestral tetrapod version of RTN4IP1 (**Figure S4A**) and expressed and purified the protein (**Figure S4B**). The enzyme was soluble and catalytically active, displaying NAD(P)H-dependent quinone reductase activity. Similar to L2HGDH, RTN4IP1 showed no strong substrate preference among the quinone intermediates tested in isolated biochemical assays (**Figure S4C; Table S1**). When incorporated into the reconstituted tetrapod metabolon, RTN4IP1 enhanced overall coenzyme Q production (**Figure S4D**). As observed for L2HGDH, PEGylation abolished this stimulatory effect without affecting quinone reductase activity, indicating that physical association with the metabolon is required for productive function (**Figures S4D-F**).

To determine whether RTN4IP1 and L2HGDH support the same or distinct reactions within the pathway, we dissected individual enzymatic modules of the quinone phase. We focused on the two methyltransferase reactions catalyzed by COQ3 and COQ5 (**Figure 2I**). Both enzymes have long been known to preferentially utilize reduced quinone substrates, although the mechanistic basis of this requirement remains unclear^38,39^. The COQ3-dependent step was monitored by initiating reactions from **6** in the presence of COQ7 and COQ9, whereas the COQ5-dependent step was assessed starting from **5** using a metabolon mixture lacking NAD^+^ (i.e. COQ7-COQ9 activity cannot operate while preserving the structural context of the assembly; **Figure 2I**). L2HGDH enhanced the COQ3-dependent reaction, with a weaker effect on COQ5. In contrast, RTN4IP1 preferentially stimulated the COQ5-dependent methylation step, while displaying a weaker effect on COQ3 (**Figure 2J**). Together, these findings revealed a degree of functional complementation between the two reductases because within the metabolon they appear to preferentially support distinct methyltransferase reactions, with L2HGDH acting primarily on the COQ3-dependent step and RTN4IP1 preferentially sustaining COQ5. However, simultaneous addition of the two reductases did not further increase coenzyme Q production beyond the effect observed for each enzyme individually implying functional compensation between the two enzymes (**Figure S4G**).

### L2HGDH forms a stable complex with the COQ3-COQ6 module of the COQ metabolon

We next investigated whether L2HGDH stably associates with the COQ metabolon. Size-exclusion chromatography showed that L2HGDH co-elutes with the reconstructed metabolon core, composed of ancestral tetrapod COQ3, COQ4, COQ5, COQ6, COQ7 and COQ9, validating its physical integration into the complex and consistent with the functional data (**Figure S5A**). Because L2HGDH preferentially enhances the COQ3-dependent step of coenzyme Q biosynthesis, we then asked whether this functional preference reflects selective interactions with defined metabolon components. Indeed, among all COQ proteins, COQ3 was the only component showing significant coevolutionary coupling with L2HGDH (**Figures 1B-C**), while independent crosslinking-MS dataset further suggested physical interactions involving both COQ3 and COQ6^35^. Analytical size-exclusion chromatography of the individual proteins (COQ3, COQ6 and L2HGDH) showed elution profiles consistent with oligomeric states larger than their expected monomers (**Figure S5B**). When COQ3 and COQ6 were mixed, the elution peak shifted toward lower retention volumes relative to the individual proteins, consistent with formation of the previously reported COQ3-COQ6 complex. Addition of L2HGDH to this binary assembly produced a sharper peak eluting at a similar volume, suggesting incorporation of a third component in the complex (**Figure S5C**). Comparison with molecular-weight standards yielded an estimated molecular mass of approximately 260 kDa, nearly twice the combined molecular weight of the three proteins (∼125 kDa), indicating the formation of a higher-order assembly containing six subunits. SDS-PAGE analysis confirmed the co-elution of COQ3, COQ6, and L2HGDH within this peak fraction, which was subsequently selected for cryo-electron microscopy (**Figure S5D**). After testing multiple vitrification conditions, only graphene oxide-coated grids yielded particles with suitable distribution and contrast within the holes (**Figure S6**). These grids produced high-quality two-dimensional class averages displaying a characteristic bee-like morphology (**Figure S7A**). Reconstruction of the three-dimensional density at 2.9 Å resolution revealed an assembly composed of two COQ3-COQ6-L2HGDH trimers arranged as a dimer of trimers (**Figure S7B**; **Table S2**). The two trimers were not equivalent: one was well resolved whereas the second appeared substantially less defined, indicating structural heterogeneity and a flexible association between the two heterotrimeric assemblies. Consistently, increasing the contour level progressively ablated density corresponding to the second trimer, while the first remained clearly visible (**Figure 3A**). Model building and refinement were therefore restricted to the well-resolved trimer.

**Figure 3.**
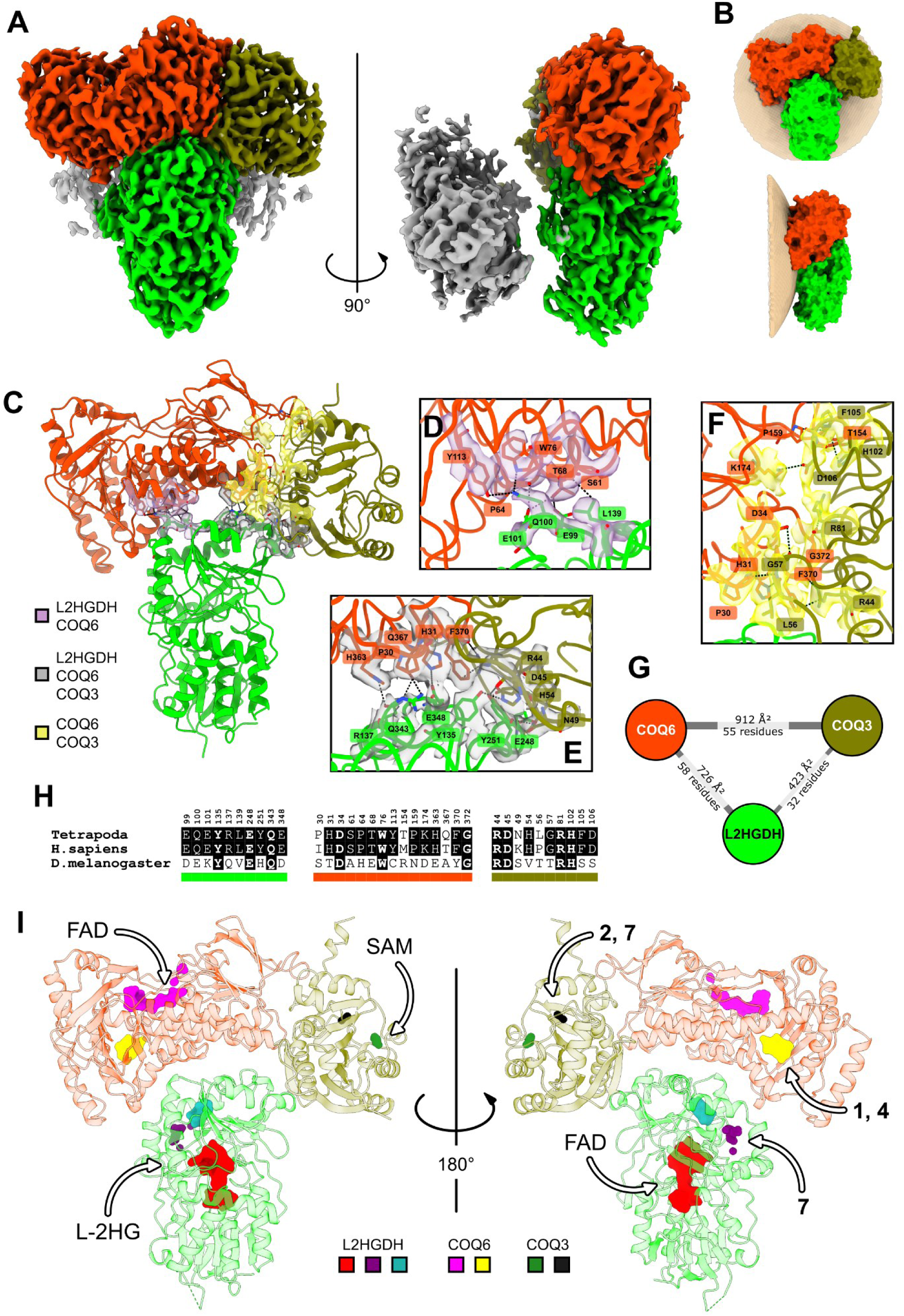
Structural integration of L2HGDH within the COQ metabolon. (A) Cryo-EM map at contour level 5 σ showing that one trimer (colored) is better resolved than the second (gray), indicating structural heterogeneity. (B) OPM prediction of the complex on the inner mitochondrial membrane, showing that the interface between the two trimers corresponds to the membrane-facing surface. (C) Cartoon representation of the COQ3-COQ6-L2HGDH trimer with interface densities shown. The different contact regions are colored according to the legend. (**D-F**) Cartoon representation of the interacting subunits with interface residues shown as sticks and the corresponding cryo-EM density displayed as a surface. Shown are the interfaces between COQ6 and L2HGDH (**D**), L2HGDH and the COQ3-COQ6 heterodimer (**E**), and COQ3 and COQ6 (**F**). Interacting residues are labeled, numbered, and colored according to the corresponding protein. (G) Schematic summary of PDBePISA analysis showing the contact surface areas between subunits together with the number of residues involved in each interface. The COQ3-COQ6 interaction forms the largest interface within the assembly, whereas L2HGDH contacts both proteins through smaller but structurally well-defined interaction regions. (H) Multiple sequence alignment excerpts of L2HGDH, COQ6 and COQ3 orthologs, showing only regions containing residues involved in the COQ3-COQ6-L2HGDH interfaces. Sequences from ancestral tetrapod, human and *Drosophila* orthologs are shown. Interface residues are indicated, and conserved residues are highlighted with black boxes. (I) Cartoon representation of the trimer highlighting the cavities identified by P2Rank and colored according to the legend. Arrows indicate the molecules predicted to occupy each cavity based on Boltz2 co-folding simulations or directly observed in the electron density.

### Structural organization of the COQ3-COQ6-L2HGDH assembly

The interface between the two trimers observed in the hexameric reconstruction is predominantly formed by hydrophobic surfaces. Density associated with this region was less well resolved, particularly for portions of COQ3 and L2HGDH involved in inter-trimer contacts (**Figure S8**). Orientation of the trimer using OPM^41^ predicts that this surface faces the inner mitochondrial membrane, supporting the idea that the observed hexamer arises from association of membrane-facing hydrophobic regions in the absence of a lipid bilayer (**Figure 3B**). Examination of the refined trimer revealed an interaction network in which COQ3 and COQ6 form the central interaction module, while L2HGDH associates with both proteins simultaneously (**Figure 3C-G**). The most extensive interface is formed between COQ3 and COQ6 (912 Å^2^), as indicated by PISA analysis (**Figures 3F-G**). L2HGDH is positioned along one side of this heterodimer and establishes two major interaction regions. The first involves a surface loop encompassing residues 99-101 that contacts COQ6 (**Figure 3D**), whereas the second is located directly at the COQ3-COQ6 interface (**Figure 3E**). In this latter region, residues such as Phe370 and His31 of COQ6 participate in interactions with both COQ3 and L2HGDH, effectively bridging the three proteins within a single assembly. Remarkably, cryo-EM analysis of the binary COQ3-COQ6 complex in the absence of L2HGDH revealed two distinct tetrameric assemblies with comparable populations, both built from the same COQ3-COQ6 heterodimer but differing in the relative orientation of the two dimers (head-to-head and head-to-tail, respectively; **Figure S9**; **Table S2**). The heterodimeric COQ3-COQ6 interface is preserved in both binary reconstructions, whereas the contacts mediating dimer-of-dimer formation differ substantially, indicating that the COQ3-COQ6 heterodimer represents a stable interaction module while its higher-order assemblies are weakly constrained. The ternary COQ3-COQ6-L2HGDH structure extends this module by positioning L2HGDH against the COQ3-COQ6 interface, fully consistent with contacts identified by mining a previously published large-scale crosslinking-MS dataset (**Figure S10**). Notably, most residues directly involved in the COQ3-COQ6-L2HGDH interface are conserved between the ancestral tetrapod and human proteins but are less conserved in more distant metazoan orthologs such as *Drosophila* (**Figure 3H**). This pattern suggests that the assembly may represent a tetrapod-conserved feature of the COQ metabolon rather than a broadly conserved metazoan interaction.

### Active-site architecture of COQ3, COQ6 and L2HGDH

To gain mechanistic insight into substrate recognition and catalysis, we analyzed the active-site architecture of the three enzymes combining the cryo-EM structure with cavity analysis, ligand co-folding and crystallographic data. COQ6, an aromatic hydroxylase catalyzing two reactions in coenzyme Q (**Figure 1A**), contains a large cavity exposed toward the soluble face of the assembly and corresponding to the expected FAD-binding region (**Figure 3I**; magenta). Neither cryo-EM density (**Figure 4A**) nor UV-visible spectroscopy (**Figure 4B**) revealed bound FAD, consistent with the observations that recombinant COQ6 is purified predominantly in its apo state^10^. To investigate substrate binding, we performed Boltz2 co-folding modelling with FAD and intermediate **4** (**Figure 1A**). Comparison between the predicted complex and the experimental structure suggests that ligand binding is accompanied by a conformational rearrangement of a loop that opens communication between the FAD and substrate cavities, whereas in the apo structure the two regions remain separated (**Figures 4C-D**). As a consequence, the substrate’s aromatic ring can bind deep inside the protein with its carbon atom undergoing hydroxylation being adjacent to the reactive N5-C4a locus of the flavin (**Figures 4E-F**). The flavin’s dimethylbenzene ring remains partially solvent exposed and, in this position, it can readily receive electrons from an external donor such as a ferredoxin (FDX2; **Figure 1A**). In summary, this arrangement satisfies all the geometric and structural requirements expected for the flavin-dependent hydroxylation.

**Figure 4.**
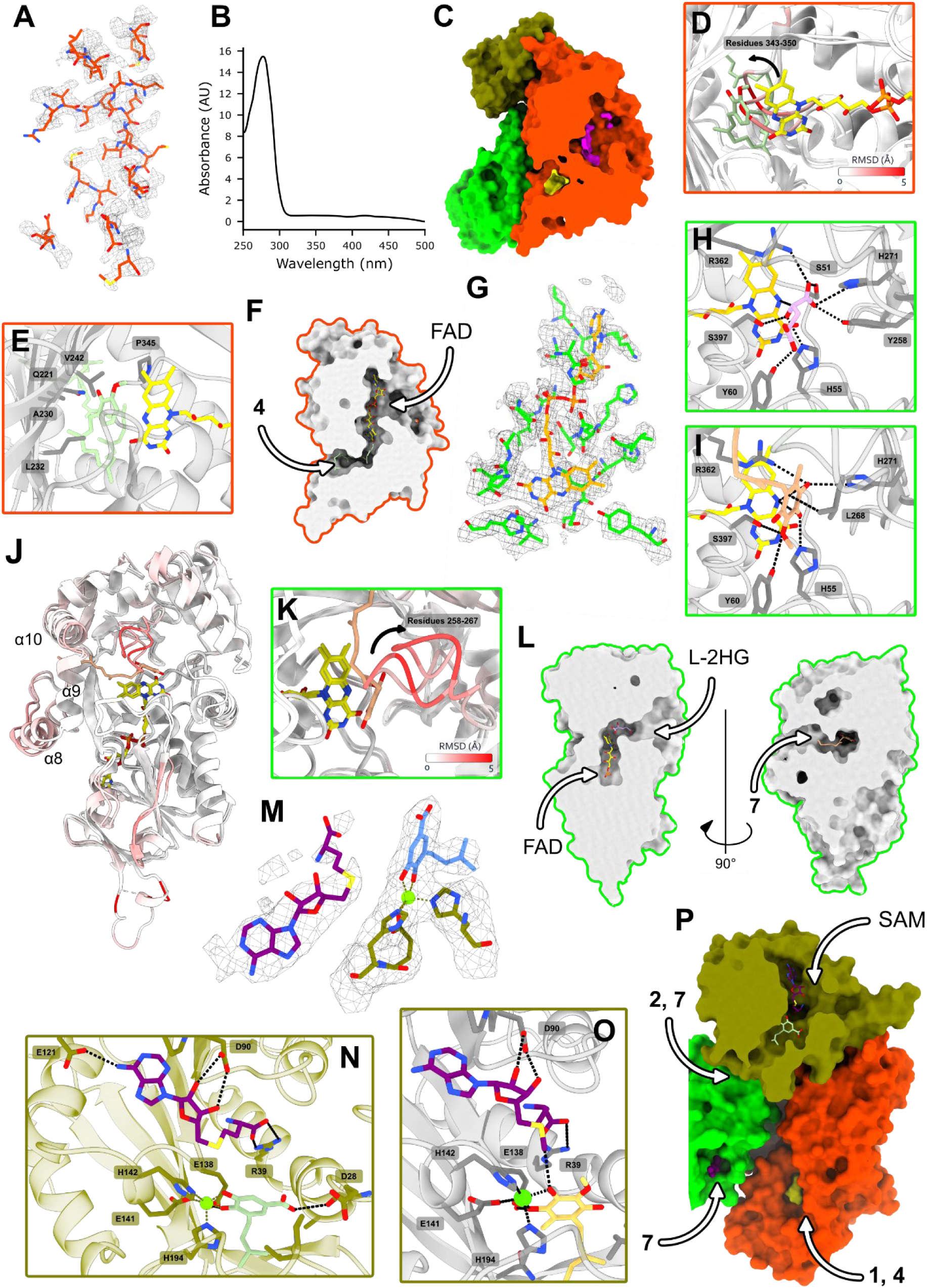
Catalytic architecture of the COQ3-COQ6-L2HGDH module. (A) Electron density of COQ6 active site. (B) UV-visible absorption spectrum of COQ6 (250 μM), showing very low signal of the characteristic spectrum of oxidized FAD. (C) Surface section of the experimentally determined COQ3-COQ6-L2HGDH trimer. In the cryo-EM structure, the FAD-binding cavity and substrate-binding cavity are separated as highlighted by the two distinct pockets identified by P2Rank analysis (yellow and magenta). (D) Cartoon representation of the superposition between the Boltz2 prediction and the experimental COQ6 structure, colored according to Cα RMSD values as indicated by the scale bar. A flexible loop (residues 343-350) is highlighted that undergoes a conformational rearrangement in the ligand-bound Boltz2 model. Movement of this loop creates space for both FAD and **4** and establishes communication between the cofactor-and substrate-binding cavities, suggesting a catalytically competent conformation. (E) Detailed view of Boltz2 co-folding models containing FAD and **4** (F) Surface section of the Boltz2 co-folding model of COQ6 with FAD (yellow) and **4** (green) shown as sticks. (G) Electron density of L2HGDH active site (contour level 4 σ). (**H-I**) Cartoon representations of Boltz2 co-folding models containing FAD and L-2-hydroxyglutarate (**H**) or FAD and **7** (**I**). Possible H-bond interactions are indicated by dashed lines. Inclusion of L-2-hydroxyglutarate prevents simultaneous accommodation of **7** in the catalytic region adjacent to FAD. (J) Superposition of the Boltz2 model and the experimental L2HGDH structure, colored according to Cα RMSD values as indicated by the scale bar. The three hydrophobic α-helices forming the characteristic horseshoe-shaped cavity proposed to accommodate the isoprenoid tail of **7** are labelled. (K) Enlarged view of panel **I**, highlighting a conformational rearrangement of residues 258-267. In the ligand-bound Boltz2 model, movement of this loop opens communication between the quinone-access cavity and the FAD active site. (L) Surface section of the Boltz2 co-folding prediction of L2HGDH with FAD (yellow) and L-2-hydroxyglutarate (lilac, left) or **7** (salmon, right) shown as sticks. Arrows indicate the predicted access routes of the quinone substrate and L-2-hydroxyglutarate. (M) 2Fo-Fc electron density of the 3.2 Å resolution crystal structure of COQ3 active site bound to S-adenosyl-homocysteine and **2**. The contour level is 1.2 σ. (N) Detailed view of the COQ3 active site shown as a cartoon with S-adenosyl-homocysteine and **2** displayed as sticks. The magnesium ion and coordinating residues are shown explicitly. Possible H-bond interactions are indicated by dashed lines. (O) Cartoon representation of the Boltz2 co-folding model containing SAM (purple) and reduced **7** (yellow). The two ortho hydroxyl groups of **7** coordinate the catalytic magnesium ion, while the hydroxyl group targeted for methylation is oriented toward the SAM methyl donor, generating a geometry compatible with catalysis. (P) Surface section of the experimentally determined COQ3-COQ6-L2HGDH trimer. Arrows indicate the predicted substrate access routes; SAM, **2** and **7** for COQ3, **1** and **4** for COQ6, **7** (and possibly **5)** for L2HGDH (Figure 1A). The SAM-binding site is exposed toward the solvent-accessible face of the complex, whereas the substrate-binding tunnel opens toward the hydrophobic surface shared with COQ6 and L2HGDH.

L2HGDH displayed well-defined density for the tightly bound FAD (**Figure 4G**). Cavity analysis identified two major pockets (**Figures 3I**). The first corresponds to the canonical FAD-dependent active site and extends toward the solvent-accessible surface, whereas the second cavity is located within a bundle of three α-helices (α8-10) forming a horseshoe-shaped structure on the hydrophobic face of the complex. Co-folding simulations with FAD and L-2-hydroxyglutarate predicted that the substrate occupies the cavity in direct contact with the flavin (**Figures 4H**). When **7** was included in the prediction, no space was available to accommodate both substrates simultaneously (**Figure 4I**). Indeed, **7** occupies the same catalytic region as the L-2-hydroxyglutarate with its isoprenoid tail extended towards the three-helix bundle (**Figure 4J**). Notably, a conformational rearrangement of a flexible loop is predicted to establish communication between the quinone-access tunnel and the FAD site (**Figure 4K**). These structural data features are consistent with a mechanistic model whereby L-2-hydroxyglutarate and the quinone electron acceptor alternatively bind to the same active-site cavity in front of the flavin, in agreement with ping-pong kinetics (**Figure 2E**). However, even if the binding cavity is the same, the two substrates may follow different routes for accessing the binding site. The quinone substrate may enter through a tunnel leading from the membrane surface whereas L-2-hydroxyglutarate may follow a different path that allows diffusion from the water-exposed surface region of the protein (**Figure 4L**).

COQ3 was the least well-resolved component of the cryo-EM reconstruction, particularly in the region facing the second, less-defined trimer, where part of the polypeptide chain could not be confidently modeled (**Figure S8A**). To obtain additional structural information, we solved a crystal structure of ancestral tetrapod COQ3 at 3.2 Å resolution, (**Figure 4M** and **Table S3**). The structure obtained in complex with S-adenosyl-homocysteine and **2** revealed a monoatomic ligand, likely to be magnesium ion coordinated by His142, Glu141, and His194 (**Figure 4N**). Consistently, Boltz2 predictions with **7** showed that the two substrate’s ortho hydroxyl groups coordinate this magnesium while positioning the reactive hydroxyl group toward the methyl donor of SAM, generating a geometry compatible with catalysis (**Figure 4O**). This arrangement provides a structural explanation for the requirement of the aromatic ring being in the reduced catechol state during methylation. Superposition of the crystal structure onto the cryo-EM trimer further revealed that the SAM-binding pocket is exposed toward the solvent-accessible face of the assembly, whereas the prenylated methyl-accepting substrates can reach the catalytic center through a tunnel opening toward the hydrophobic surface of the metabolon. Notably, this tunnel is located on the same membrane-facing side used for substrate access by both COQ6 and L2HGDH, suggesting the existence of a shared hydrophobic reaction surface that facilitates the channeling of isoprenoid-containing intermediates between neighboring enzymes (**Figure 4P**).

### L-2-hydroxyglutarate accumulates in COQ4 deficient patients

The identification of L2HGDH as a functional component of the COQ metabolon prompted us to revisit a previously reported metabolic feature of COQ4 deficiency. Earlier studies described the accumulation of 2-hydroxyglutarate in biological samples from COQ4-deficient patients, although the relative contribution of the D-and L-enantiomers was not determined ^42^. To clarify this relationship, we quantified both L-and D-2-hydroxyglutarate by UPLC-MS/MS in urine samples from two pediatric individuals carrying pathogenic COQ4 variants (c.[437T>G], p.[Phe146Cys], homozygous; c.[469C>A];c.[718C>T]; p.[Gln157Lys],p.[Arg240Cys], compound heterozygous), together with samples from healthy controls and patients affected by L-2-or D-2-hydroxyglutaric aciduria, which served as positive controls (**Figure 5A**). Patients with L-2-and D-2-hydroxyglutaric aciduria displayed marked and stereospecific accumulations of the corresponding metabolite. Remarkably, urine samples from both COQ4-deficient individuals also exhibited elevated concentrations of L-2-hydroxyglutarate compared with healthy controls (**Figure 5A**). Increased levels of D-2-hydroxyglutarate were likewise detected, although with lower values than those observed for the L-enantiomer. To determine whether this metabolic signature was also present at the cellular level, we analyzed primary skin fibroblasts derived from three COQ4-deficient patients (c.[155T>C];c.[521_523delCCA], p.[Leu52Ser];p.[Thr174del], compound heterozygous; c.[458C>T], p.[Ala153Val], homozygous; c.[437T>G], p.[Phe146Cys], homozygous). Consistent with the urine measurements, COQ4-deficient fibroblasts displayed significantly higher levels of L-2-hydroxyglutarate relative to control cells (**Figure 5B**). In contrast, D-2-hydroxyglutarate levels showed no or only partial increase (**Figure 5C**). Together, these findings establish that impairment of coenzyme Q biosynthesis is associated with altered hydroxyglutarate homeostasis *in vivo* and in patient-derived cells, supporting the pathophysiological relevance of the biochemical connection between L-2-hydroxyglutarate metabolism and coenzyme Q biosynthesis. Alterations of L-2-hydroxyglutarate levels might thereby contribute to the clinical symptoms of coenzyme Q deficiency and should be evaluated in coenzyme Q deficient patients.

**Figure 5.**
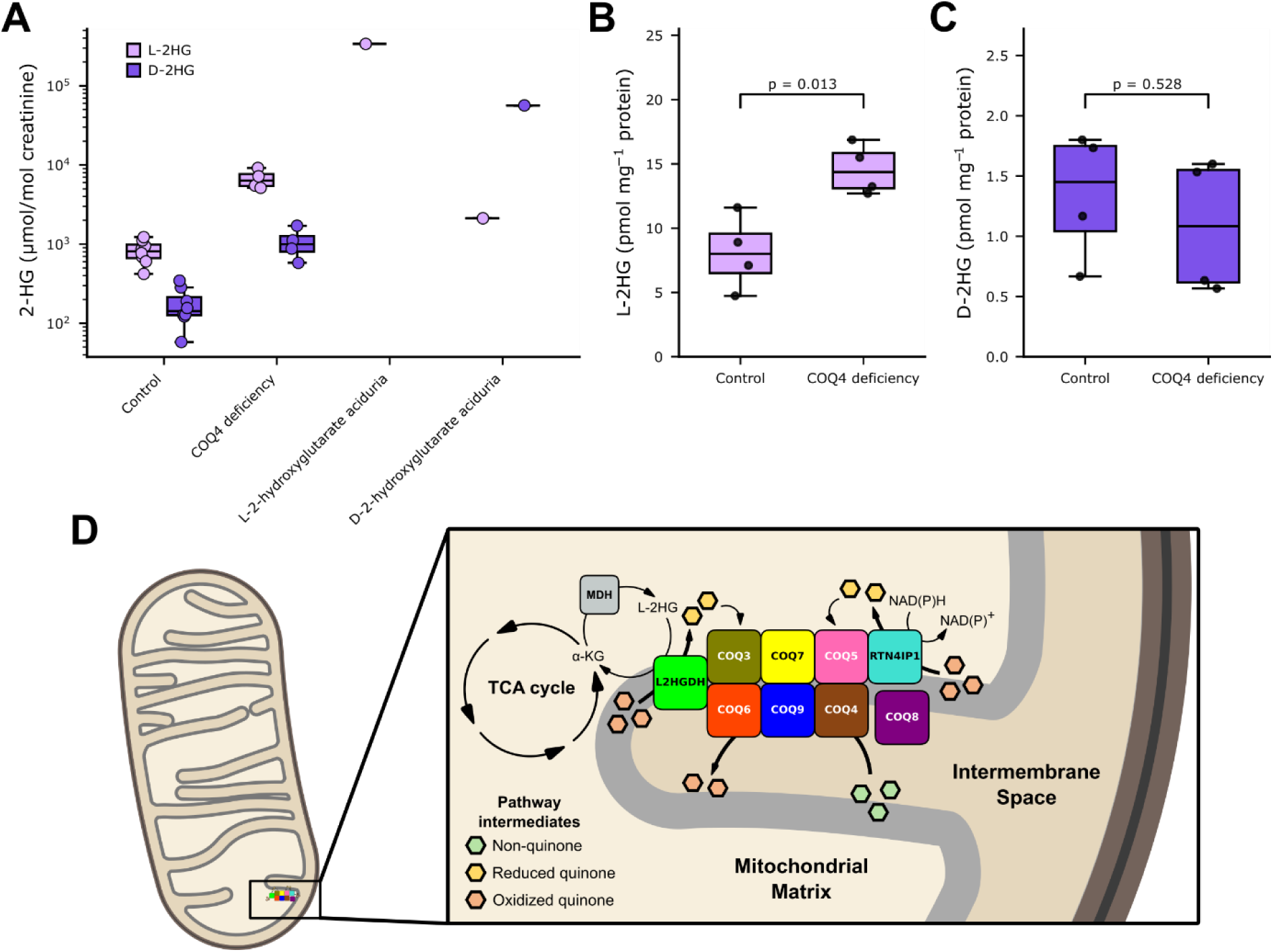
L-2-hydroxyglutarate accumulation in COQ4 deficiency supports a physiological link between L2HGDH and the COQ metabolon (**A**) Measurements of L-2 and D-2 hydroxyglutarate levels in urine samples of children with genetically confirmed COQ4 deficiency (n = 4; pooled results of 2 independent individuals) compared to healthy controls (n = 8, pooled results) and individuals with genetically confirmed L-2-and D-2-hydroxyglutarate aciduria (n = 1, respectively; “positive controls”). (**B-C**) Measurements of L-2-hydroglutarate (**B**) and D-2-hydroglutarate (**C**) levels in cultured primary skin fibroblasts (4 independent experiments) from three patients with COQ4 deficiency (3 cell lines, pooled results) compared to age and sex matched controls (3 cell lines, pooled results). (D) Functional coupling between L2HGDH and the COQ metabolon in the mitochondria. The methyltransferase enzymes COQ5 and COQ3 are supported by the local quinone reductase activities of RTN4IP1 and L2HGDH, respectively. L-2-hydroxyglutarate (L-2HG), produced by the promiscuous activity of malate dehydrogenase (MDH), is oxidized to α-ketoglutarate (α-KG) by L2HGDH while electrons are transferred to reduce quinone intermediates (yellow) of the coenzyme Q biosynthetic pathway. COQ4 defects may therefore lead to L-2HG accumulation by limiting the availability of oxidizable quinone intermediates as electron acceptors.

## Discussion

L-2-hydroxyglutarate is generated primarily under hypoxic conditions^30,31^ by the promiscuous activity of enzymes such as malate and lactate dehydrogenases^43^. Recent data have demonstrated the Janus-faced nature of this metabolite that is both a signaling molecule as well as a potentially toxic compound^19^. L-2-hydroxyglutarate inhibits a broad class of α-ketoglutarate-dependent enzymes, including histone^44^ and DNA demethylases^45^ as well as DNA repair enzymes^46^, to a greater extent than its enantiomer D-2-hydroxyglutarate, a powerful oncometabolite^45^. L-2-hydroxyglutarate accumulates as an effect of loss-of-function mutations in L2HGDH, a severe disease that causes L-2-hydroxyglutaric aciduria and predispose affected individuals to neurological disorders and brain tumors^47–49^. Here, we show that L2HGDH does not simply function in the scavenging and detoxification of L-2-hydroxyglutarate. Rather it is an integral component of the coenzyme Q biosynthetic machinery, and our data indicates that this association may also have clinical implications in conditions of altered coenzyme Q and L-2-hydroxyglutarate/α-ketoglutarate metabolism. Structurally, L2HGDH associates with the COQ metabolon through COQ3 and COQ6, the dual-step enzymes of the pathway. Notably, COQ3 was the only metabolon component displaying significant coevolutionary coupling with L2HGDH (**Figure 1B**) and was previously reported to be downregulated in patients with high levels of 2-hydroxyglutarate in acute myeloid leukemia^50^, providing independent support for a functional relationship between the two proteins. Functionally, our data indicate that L2HGDH boosts coenzyme Q generation by safeguarding the catalytically favorable reduced state of the substrates of COQ3 and COQ5, the two methyltransferases necessary for coenzyme biosynthesis. Although L2HGDH can reduce multiple quinone intermediates, it is functionally biased toward the COQ3-catalysed final step of the pathway, whereas RTN4IP1, an NADPH-dependent oxidoreductase, preferentially supports COQ5 activity. This reveals a division of labor in which distinct reductases sustain specific steps of coenzyme Q biosynthesis (**Figures 5D** and **S11**).

After 50 years from its discovery^51^, our data expands the role of L2HGDH, now highlighted as a central node of mitochondrial metabolism. Its oxidoreductase activity links the homeostasis of two key cofactors: α-ketoglutarate, regenerated from its waste metabolite L-2-hydroxyglutarate, and coenzyme Q, whose biosynthesis is boosted through the localized reduction of pathway intermediates. Because L-2-hydroxyglutarate production increases under conditions of elevated mitochondrial NADH/NAD⁺ ratios, recruitment of L2HGDH into the metabolon may operate as a floodgate linking cellular redox state to coenzyme Q biosynthesis. Under conditions of redox imbalance, altered levels of L-2-hydroxyglutarate may affect the biosynthetic flux and thereby modulate coenzyme Q levels. In this framework, a fundamental takeaway of this work is that metabolons are not standalone, autonomous machines. Rather, their transient and non-stoichiometric nature allows them to structurally and functionally integrate with other systems, bridging metabolic, signaling, and regulatory pathways.

### Limitations of the study

While the L-2-hydroxyglutarate levels measured in coenzyme Q-deficient patients validate the physiological relevance of our findings, a limitation is that our biochemical and structural studies were conducted using ancestral proteins. Although these proteins share high sequence identity (>80-90%) with their human counterparts, their exact kinetic and biophysical properties may vary. Consequently, the magnitude of the effects exerted by human L2HGDH on coenzyme Q biosynthesis might differ slightly from the values reported here. Future metabolic flux studies are required to fully evaluate how L2HGDH activity and L-2-hydroxyglutarate levels influence coenzyme Q biosynthesis across diverse tissues and pathophysiological conditions, a new research avenue opened by this work.

## Methods

### Materials

All chemicals and enzymes for assays have been purchased from Sigma-Aldrich unless specified. Detergents were purchased from Anatrace. 4-Hydroxy-3-(3-methylbut-2-en-1-yl)benzoic acid (**1**) was purchased from BLD Pharmaceutics. 3,4-Dihydroxy-5-(3-methylbut-2-en-1-yl)benzoic acid (**2**), 4-hydroxy-3-methoxy-5-(3-methylbut-2-en-1-yl)benzoic acid (**3**), 2-methoxy-6-(3-methylbut-2-en-1-yl)phenol (**4**), 2-methoxy-6-(3-methylbut-2-en-1-yl)benzene-1,4-diol (**5**), 5-methoxy-2-methyl-3-(3-methylbut-2-en-1-yl)cyclohexa-2,5-diene-1,4-dione (**6**) and Sorbicillin were obtained from WuXi App Tech. Purity and ^1^H NMR of intermediates 1, 2, 3, 4, 5, 6 have been previously reported ^10,40^. UHPLC quality solvents were purchased from Romil.

### Construction of the functional interaction COQ network

To construct the functional interaction network, the complete human proteome was obtained from UniProt and used as the reference dataset. For each protein query, four independent categories of functional interaction were evaluated. Coexpression was retrieved from COXPRESdb and the Human Protein Atlas. In COXPRESdb, interactions were considered positive when the candidate gene appeared within the top 50 positions of the coexpression ranking for a given query. In the Human Protein Atlas, genes were considered coexpressed when they were assigned to the same expression cluster in at least one dataset category (Blood, Tissue, Brain, Cell line, or Single cell). Colocalization was retrieved from the ComPPI and BioGRID databases, which directly provide experimentally supported or curated protein-protein interaction hits for each protein. Coevolutionary relationships were inferred using the program *cotr* (https://github.com/lab83bio/Cotransitions), employing three versions of the OrthoDB database (v10.1, v11.0, and v12.1) at the Eukaryota level (at2759). For OrthoDB v12.1, additional analyses were performed at three additional taxonomic levels: Metazoa (at33208), Vertebrata (at7742), and Mammalia (at40674). Candidate interactions were considered significant when the cotr analysis returned a *p-value* < 1e-3 for a given query. Evidence for domain fusion was obtained from the Pfam database, where interactions were considered positive when two domains were found fused within at least one annotated domain architecture. All analyses were automated using a custom Jupyter notebook (*network.ipynb*), provided in the **Source Data**. The resulting table (*final.csv*) contains, for each pair of proteins, the query, the candidate interactor, and the data supporting their functional linkage. The dataset was subsequently filtered to retain only interactions involving genes encoding core components of the COQ metabolon (COQ3, COQ4, COQ5, COQ6, COQ7, and COQ9). For each candidate gene, an interaction score was calculated by counting the number of hits supporting its association with any COQ component (**Figure 1B**). Candidates with a score ≥10 were retained for downstream analysis and visualization (**Figure 1C**).

### Ancestral sequence reconstruction

Ancestral sequences were reconstructed using the ASR.ipynb notebook provided in the **Source Data**. Briefly, L2HGDH and RTN4IP1 orthogroups were retrieved at the Vertebrata level from OrthoDB v12.2 (1540610at7742 and 191779at7742, respectively). Sequences were aligned using MAFFT v7, followed by iterative cleaning and trimming steps implemented in the ASR.ipynb workflow. The curated alignments were then used to reconstruct phylogenetic trees with RAxML-NG v1.2.2 using the JTT+G4 substitution model and 100 bootstrap replicates, with species-tree constraints applied during reconstruction (385 sequences for L2HGDH and 262 sequences for RTN4IP1). Ancestral sequences for each internal node were inferred using the *--asr* option of the same software. All input files, intermediate steps, and final outputs are available in the **Source Data**.

### Protein expression and purification

Protocols for the expression and purification of the COQ metabolon components (COQ3, COQ4, COQ5, COQ6, COQ7, COQ8B, COQ9, FDXR, and FDX2) have been previously described^10^. Plasmids for expression of N-terminally His-SUMO-tagged L2HGDH (pET-28) and RTN4IP1 (pET-24) were obtained from Genscript. The mitochondrial targeting sequences were removed based on Mitofates predictions^52^. Plasmids were transformed into *E. coli* BL21 CodonPlus cells. Cultures were grown at 37 °C to an OD_600_ of 0.6 and protein expression was induced with 0.2 mM IPTG overnight at 20 °C. Cell pellets were resuspended in Lysis Buffer (20 mM potassium phosphate, pH 7.8, 300 mM NaCl, 1% Triton X-100, 10% glycerol, DNase, pepstatin) and incubated for 20 minutes while stirring. Cells were then lysed by sonication (1 s on / 1 s off cycles, 15 minutes total on-time, 30% amplitude). The lysate was clarified by centrifugation at 56,000 × g for 45 minutes, and the supernatant was loaded onto an ÄKTA Pure system equipped with a 5 mL HisTrap FF column equilibrated in Loading Buffer (50 mM HEPES pH 7.5, 250 mM NaCl, 0.018% DDM). The column was washed with 5 column volumes (CV) of loading buffer supplemented with 25 mM imidazole. Proteins were eluted using a linear gradient over 7 CV from 25 to 500 mM imidazole in the same buffer. Fractions were analyzed by SDS-PAGE, pooled, and desalted using a 5 mL HiTrap desalting column to remove imidazole and exchange into Loading Buffer. The His-SUMO tag was removed by overnight digestion at 4 °C using SUMO protease (1% v/v). The cleaved protein was separated from the tag by a second HisTrap purification step under the same buffer conditions, collecting the protein in the flow-through. The final purification step consisted of size-exclusion chromatography using a preparative Superdex 200 Increase 10/300 GL column equilibrated in 25 mM HEPES pH 7.5 and 50 mM NaCl for L2HGDH. RTN4IP1 was purified using the same buffer conditions supplemented with 0.018% DDM. Purified proteins were flash-frozen in liquid nitrogen and stored at-80 °C until further use.

### Kinetic characterization of L2HGDH, RTN4IP1, and FSP1

Purified proteins were used to determine kinetic parameters in the presence of coenzyme Q and other quinone substrates. For L2HGDH activity, we established a coupled assay with glutamate dehydrogenase. The reaction mixture contained 20 mM HEPES pH 7.5, 0.075 μM L2HGDH, 5 mM NH_4_Cl, 350 μM NADH, 1μM GDH and variable concentrations of the quinone substrates tested. Reactions were initiated by addition of 2 mM L-2-hydroxyglutarate. For RTN4IP1, the same buffer was used, with enzyme at a final concentration of 0.164 μM. Activity was monitored directly by following NADH consumption at 340 nm. All measurements were performed on an Agilent Cary 60 spectrophotometer at 37 °C using a quartz cuvette with a 1 cm path length. FSP1 assays were performed using purified protein and the previously published assay conditions^53^.

### Protein PEGylation

PEGylated L2HGDH and RTN4IP1 were obtained by following protocols previously described^40^. Briefly, 2 mg ml-1 O-methyl-O’-succinyl polyethylene glycol 5,000 N-succinimidyl ester (methoxyPEG-NHS) were mixed with 1 mg ml-1 protein in 10 mM sodium bicarbonate. Reaction was carried out for one hour at room temperature. Reaction was quenched with 100 mM Tris-HCl pH 8 at 4 °C. The excess of reactant was removed by desalting using a HiTrap column (Cytiva) pre-equilibrated in 25 mM HEPES pH 7.5 and 50 mM NaCl and an Äkta Pure system (Cytiva).

### Small scale reactions

Small scale reactions were carried out overnight at 25°C 400 rpm as previously reported ^40^. Briefly, COQ3-9 were used at a 1 μM final concentration with 2 μM FDXR and FDX2, 250 μM FAD, 1 mM S-adenosylmethionine, 150 μM MgCl₂, 25 μM ZnCl₂, 2 mM L-2-hydroxyglutarate, 1 mM of the starting substrate, NAD(P)H and ATP regeneration systems. NAD(P)H regeneration consisted of 300 μM NADP⁺ and NAD⁺, 1.2 U glucose dehydrogenase, and 1.2 mM glucose. ATP regeneration consisted of 1 mM ADP, 4 U pyruvate kinase, 5 mM inorganic phosphate, and 5 mM phosphoenolpyruvate. Reactions were quenched by adding 33% (v/v) final acetonitrile. For identification of coenzyme Q and its intermediates, all quinones were oxidized by adding 10 mM benzoquinone for 30 min at RT. Prior to injection in LC-MS, reactions were diluted 1:10 in 2:1 water:acetonitrile containing 0.1% (v/v) formic acid and 1 μM sorbicillin as an internal standard. For identification of L-2-hydroxyglutarate reactions were diluted 1:10 in water containing 10 mM ammonium acetate and 10 μM glutarate as an internal standard.

### LC-MS

#### Coenzyme Q1 and intermediates

Intermediates and coenzyme Q_1_ were detected and quantitated as previously described ^40^. Analysis was carried out on a X500B quadrupole-TOF (Sciex) coupled to an ExionLC (Sciex) mounted with a Kinetex EVO C18 column (100 mm × 2.1 mm, 2.6 μm particle size; Phenomenex). The flow rate was 0.2 ml min⁻¹. Mobile phases consisted of water (A) and acetonitrile (B), both containing 0.1% (v/v) formic acid. The gradient was: 2% B (0-0.1 min), 2-66% B (0.1-32.0 min), and 66-2% B (32.0-35.0 min). MS parameters were: curtain gas 30 psi, ion source gas 1 at 45 psi, ion source gas 2 at 55 psi, temperature 450 °C, spray voltage +5,500 V, declustering potential +50 V, collision energy +10 V, and full-scan range m/z 50-1,000 in positive mode. Mass calibration was performed prior to analysis using ESI positive calibration solution (Sciex). Analytes identification was performed with an allowed mass error of ±5 ppm, peaks integration was performed on the [M+H]^+^ extracted ions chromatograms and quantitation was performed on external calibrations using sorbicillin as an internal standard. Data processing was carried out with Analytics Software within Sciex OS (Sciex).

#### L-2-hydroxyglutarate

Analysis was carried out on a X500B quadrupole-TOF (Sciex) coupled to an ExionLC (Sciex) mounted with a Kinetex HILIC 100 Å column (100 mm × 2.1 mm, 2.6 μm particle size; Phenomenex). The flow rate was 0.25 ml min⁻¹. Mobile phases consisted of water (A) and acetonitrile (B), both containing 10 mM ammonium acetate. The gradient was: 95-90% B (0-1.5 min), 90-15% B (1.5-9 min), 15% B (9-12 min), 15-95% B (12-12.5 min) and 95% B (12.5-20 min). MS parameters were: curtain gas 30 psi, ion source gas 1 at 45 psi, ion source gas 2 at 55 psi, temperature 450 °C, spray voltage-4,500 V, declustering potential-80 V, collision energy-10 V, and full-scan range m/z 50-300 in negative mode. Mass calibration was performed prior to analysis using ESI negative calibration solution (Sciex). Analytes identification was performed with an allowed mass error of ±5 ppm, peaks integration was performed on the [M-H]^-^ extracted ions chromatograms and quantitation was performed on external calibrations using glutarate as an internal standard. Data processing was carried out with Analytics Software within Sciex OS (Sciex).

#### CryoEM grids preparation

Graphene oxide-coated grids for cryo-EM were prepared in-house following a previously published protocol^54^. Quantifoil R 1.2/1.3 Cu 300 grids (Quantifoil Micro Tools GmbH, Germany) were glow-discharged for 60 s at 30 mA, in a PELCO easyGlow system (Ted Pella, USA). A 10 μL drop of graphene oxide solution (0.02 mg mL-1 grOX supplemented with 0.01% DDM) was then applied to each grid. Coated grids were prepared one day prior to vitrification. For sample preparation, protein was applied at a final concentration of 2 μM. Grids were vitrified using a Vitrobot Mark IV instrument (Thermo Fisher Scientific, USA) operated at 4 °C and 95% humidity. A total sample volume of 4 μL was applied to each grid, followed by a waiting time of 5 s and blotting for 5.5 s before plunge-freezing in liquid ethane.

### Cryo-EM data collection and image processing

#### COQ3-COQ6-L2HGDH data processing

Cryo-EM data were collected at the CM01 beamline (ESRF, Grenoble) using a Titan Krios transmission electron microscope equipped with a Gatan K3 direct electron detector operated in counting mode and a Gatan Quantum LS energy filter. Data were acquired at a nominal magnification of 105,000x, corresponding to a calibrated pixel size of 0.839 Å per pixel. Two images were collected per hole over a defocus range of −0.8 to −2.2 μm with 0.2 μm step increments. The energy filter slit width was set to 20 eV, and no phase plate was used. Movies were recorded in TIFF format with LZW compression, without gain normalization, using a total exposure time of 1.93 s fractionated into 48 frames. The dose rate was 18.7 electrons per pixel per second, corresponding to a total accumulated dose of 51.15 electrons Å⁻² per movie. A total of 26,448 movies were collected. Motion correction and dose-weighting were initially performed in CryoSPARC using patch motion correction, followed by patch CTF estimation. Micrographs were filtered based on the following quality thresholds: defocus average between 0 and 30,000 Å, CTF fit up to 6.0 Å, relative ice thickness up to 1.5, and total motion below 80 pixels. After curation, 24,310 micrographs were retained for downstream processing. Initial particle picking was performed using blob picking with a particle diameter of 130 Å. Particles were extracted using a box size of 360 pixels, yielding 28,363,792 initial particle images. These particles were subjected to multiple iterative rounds of 2D classification to remove contaminants, false positives, damaged particles, and poorly aligned classes. Cleaned particle subsets were then used for ab initio reconstruction with eight initial models to capture structural heterogeneity within the dataset. The resulting classes were further refined by heterogeneous refinement, and particles corresponding to the best-resolved population were selected for final reconstruction. Non-uniform refinement was performed in C1 symmetry to generate the consensus 3D map. Final map quality was further improved using reference-based motion correction. Resolution was estimated using the gold-standard Fourier shell correlation (FSC) 0.143 criterion. Map visualization, local resolution analysis, and model fitting were carried out in CryoSPARC and UCSF ChimeraX. The pipeline of the process is shown in **Figure S6**.

#### COQ3-COQ6 data processing

Cryo-EM data were collected at the CM01 beamline (ESRF, Grenoble) using a Titan Krios transmission electron microscope equipped with a Gatan K3 direct electron detector operated in counting mode and a Gatan Quantum LS energy filter. Data was acquired at a nominal magnification of 81,000x corresponding to a calibrated pixel size of 1.06 Å per pixel. Two images were collected per hole over a defocus range of −1.0 to −2.2 μm with 0.2 μm step increments. The energy filter slit width was set to 20 eV, and no phase plate was used. Movies were recorded in TIFF format with LZW compression, without gain normalization, using a total exposure time of 1.8 s fractionated into 50 frames. The dose rate was 32 electrons per pixel per second, corresponding to a total accumulated dose of 51.4 electrons Å⁻² per movie. A total of 17,828 movies were collected. Motion correction and dose-weighting were initially performed in CryoSPARC using patch motion correction, followed by patch CTF estimation. Initial particle picking was performed using blob picking with a particle diameter of 120 Å. Particles were extracted using a box size of 256 pixels, yielding 26,300,182 initial particle images. These particles were subjected to multiple iterative rounds of 2D classification to remove contaminants, false positives, damaged particles, and poorly aligned classes. Cleaned particle stacks were then used to re-pick using Topaz which substantially improved particle picking. Ab initio reconstructions were then generated with eight initial models to capture and segregate the head-to-head and head-to-tail structural heterogeneity within the dataset. The resulting classes were further refined by heterogeneous refinement, and particles corresponding to the best-resolved population were selected for final reconstruction. Non-uniform refinement was performed in C2 symmetry to generate the consensus 3D map. Resolution was estimated using the gold-standard Fourier shell correlation (FSC) 0.143 criterion. Map visualization and local resolution analysis were carried out in CryoSPARC and UCSF ChimeraX. No models were generated based on the relatively high resolution of the models.

#### Cryo-EM model building, refinement and validation

Initial atomic models for model building were generated from Boltz2 predictions for COQ6 and L2HGDH, whereas the experimentally determined crystal structure of ancestral tetrapod COQ3 was used as the starting model for COQ3. The individual models were first rigid-body fitted into the cryo-EM density using the *Fit in Map* tool in UCSF ChimeraX. The fitted coordinates were then manually inspected and rebuilt in Coot v1.1.14. Side-chain conformations were adjusted according to the experimental density, and residues lacking interpretable density were removed from the model, including poorly resolved regions of COQ3 and one flexible loop of L2HGDH.

The model was refined iteratively using Phenix real-space refinement v2.0-5936. Refinement was performed against the final CryoSPARC sharpened consensus map using the resolution estimated from the final CryoSPARC non-uniform refinement. Global minimization, NQH flips, local grid search, and atomic displacement parameter refinement were enabled, while other parameters were kept at their default values. After each refinement round, the model was manually inspected and corrected in Coot, followed by additional cycles of real-space refinement in Phenix. The final refined model was validated using the statistics returned by the final Phenix refinement run, including model-to-map correlation, stereochemical geometry, Ramachandran statistics, rotamer outliers, clashscore, bond and angle deviations, and atomic B-factor distributions. The final refined model was validated using Phenix and MolProbity statistics as reported in **Table S2**.

#### X-ray crystallography

Crystallization was performed using 10 mg/mL COQ3 in 50 mM TRIS (pH 8.5), 10 mM NaCl, 150 µM MgCl_2_, and 1 mM COQ3 substrate. Small, transparent needle-like crystals appeared at 4°C after 3 days using the vapour-diffusion sitting-drop method from the NeXtal PEGs Suite crystal screening (Molecular Dimensions). For microseeding, small crystals grown in 0.1 M MES (pH 6.5) with 15% (w/v) PEG 20000 were crushed. The final drops for crystallization were prepared using 0.15 μL of protein, 0.05 μL of seeds, and 0.2 μL of reservoir solution. Larger crystals developed after several days in 0.1 M sodium HEPES (pH 7.5) with 15% (w/v) PEG 20000. They were harvested and cryoprotected with 30% PEG 400, supplementing the cryoprotecting solution with 1 mM COQ3 substrate (**2**) and 200 µM MgCl_2_. Data collection was carried out at the European Synchrotron Radiation Facility in Grenoble, France, and the data were processed using the XDS and CCP4 software packages. Molecular replacement was performed using an AlphaFold model and MolRep within the CCP4i suite^55^. Subsequent model building and refinement were conducted with COOT and Refmac5.

#### Computational structural modeling, pocket analysis and visualization

Ligand co-folding modelling was performed using Boltz2 (https://github.com/jwohlwend/boltz), installed locally and run on a workstation equipped with an NVIDIA GeForce RTX 3080 GPU with 12 GB RAM. Input YAML files and the scripts used to submit the modeling jobs are provided in the **Source Data**. For each protein-ligand or protein-complex system, 20 independent predictions were generated. Models were ranked according to the ipTM score, and the highest-scoring model compatible with the expected catalytic geometry was selected for structural analysis and figure preparation.

Protein cavities were identified using P2Rank (https://github.com/rdk/p2rank) with default parameters. All predicted pockets were inspected, and cavities relevant to cofactor binding, substrate access, or quinone accommodation were visualized on the experimental or predicted structures. Membrane orientation of the COQ3-COQ6-L2HGDH complex was predicted using the PPM 3.0 web server implemented in the OPM database (https://opm.phar.umich.edu/ppm_server3_cgopm). Calculations were performed using the inner mitochondrial membrane option and allowing membrane curvature.

Structural superpositions, B-factor visualization, hydrophobicity surface coloring, and figure preparation were performed in UCSF ChimeraX v1.11.1. B-factor representations were generated from refined atomic B factors and displayed as cartoon tubes colored according to B-factor values. Hydrophobicity surfaces were colored in ChimeraX using the Kyte-Doolittle hydrophobicity scale.

#### Measurement of D-/ L-2-hydroxyglutaric acid in patient samples

Both metabolites, D-and L-2-hydroxyglutarate (Sigma-Aldrich, USA) as well as their corresponding deuterated internal standards d5-D-/ d5-L-2-hydroxyglutarate (Toronto Research Chemical, Canada) were analyzed by UPLC-MS/MS. The system consisted of an UPLC I-Class (Waters, USA) coupled to a tandem mass spectrometer Xevo-TQ-XS (Waters, USA). Electrospray ionization was performed in the negative ionization mode. Analytes were separated on an Acquity UPLC HSS T3 column (50mm x 2,1 mm x 1.8 µm) by a gradient elution. The mobile phase A consisted of 10 mM ammonium acetate containing 0.1 % formic acid and the mobile phase B was acetonitrile. A sample volume of 2 µL was injected. The column temperature was set at 40 °C and the flow rate was determined as 400 µL/min. Detection of both metabolites was performed in the multiple reaction monitoring (MRM) mode. Raw files were processed for peak integration and metabolite quantification using the TargetLynx software within MassLynx v 4.2 package (Waters, USA).

For urine analysis, the following extraction procedure was chosen: 500 µL urine sample was acidified by 10 µL 1N HCl. After adding 10 µL of internal standards (1µg/mL), a volume of 1 mL of ethyl acetate, as extraction solvent, was added to the sample. The organic phase was then dried under a stream of nitrogen. The residue was dissolved in 100 µL of the DATAN (50mg/mL dissolved in acetonitrile/acetic acid; 4:1) solution. The derivatization reagent DATAN, (+)-O, Ó-diacetyl-2-tartaric anhydride (Sigma-Aldrich, USA), was applied to provide chiral separation of D-/ L-2-hydroxyglutarate. The sample was heated for 1 h at 80 °C. After cooling to room temperature, 30 µL of the derivatized sample was diluted with 60 µL distilled water before being injected into the UPLC-MS/MS system^56,57^.

For analysis of cultured cells, the compounds D-/ L-2-hydroxyglutarate were extracted from primary human skin fibroblast pellets by adding 200 µL of MeOH/H2O including 0.1 % 1 N HCl (v/v; 9:1) and 10 µL of the deuterated internal standards. After evaporation of the extraction solvent under the nitrogen stream, the dried samples were derivatized with DATAN as described above for urine analysis.

## Statistics

Statistical analyses were performed using Python code available in the **Source data**. Barplots are shown as mean of replicates together with the single point measured. Technical replicates were averaged before statistical analysis, and independent biological replicates were used as the unit of analysis. In figure 5, comparisons between two groups were performed using two-sided Welch’s t-test. P values < 0.05 were considered statistically significant.

## Ethics Statement

The institutional review boards of the Heinrich Heine University Düsseldorf (#2021-1340) approved the work involving human individuals. Participants gave informed consent to participate in the study before taking part.

## Resource availability

### Lead contacts

Further information and requests for resources and reagents should be directed to and will be fulfilled by the lead contacts, Marco Malatesta and Andrea Mattevi

### Materials availability

No new reagents were developed in this study.

### Data and code availability

The cryo-EM maps have been deposited in the EMDB under accession codes EMD-58664, EMD-58667 and EMD-58668. Coordinates derived from EMD-58664 have been deposited in the PDB under accession code 31UC. The crystal structure of COQ3 has been deposited in the PDB under accession code 31QD. Source data and analysis notebooks are available on Zenodo (10.5281/zenodo.21029243)

## Acknowledgments

This research was funded by the ERC Advanced Grant, MetaQ (no. 101094471 to A.M.), and by the Associazione Italiana per la Ricerca sul Cancro (AIRC; Investigator Grant no. 28754 to A.M. and fellowships n° 32882 in memory of Angelo e Giancarla Giardina to A.G. and n° 33008 to M.M.). We thank Centro Grandi Strumenti in Pavia for the LC-MS and cryoEM analysis. We warmly thank the European Synchrotron Radiation Facility (ESRF) for the provision of beam-time at their synchrotron radiation and cryoEM facilities and for the excellent support during data collection. We thank Alison Berezuk for her support with cryo-EM data processing and Sophia Anna Terzi for cryo-EM grid preparation.

## Author contributions

M.M.: conceptualized the study, designed and performed experiments, manuscript writing; A.G.: designed and performed biochemical experiments; C.R.N.: structural analysis of COQ3-COQ6 complex, preliminary experiments; D.H. clinical biochemistry, analysis of cells from patients; D.M. help with the biochemical experiments; D.C. experiments on FSP1, COQ3 crystal structure; A.S.; clinical biochemistry and analysis of cells from patients; F.D. supervision, writing, clinical data. A.M.: conceptualization, methodology, writing, supervision, and funding acquisition.

## Declaration of generative AI and AI-assisted technologies in the writing process

During the preparation of this work, the authors used ChatGPT to optimize the code for the bioinformatics analysis and generation of plots. After using this tool, the authors reviewed and edited the content and took full responsibility for the publication.

## Abbreviations

L2HGDH: L-2-hydroxyglutarate dehydrogenase
TCA: tricarboxylic acid
PEG: methoxy-polyethylene glycol

