## Supplementary Figures and Tables for "L-2-hydroxyglutarate recycling is linked to coenzyme Q biosynthesis"

**Content:**

Figures S1-S11

Tables S1-S3

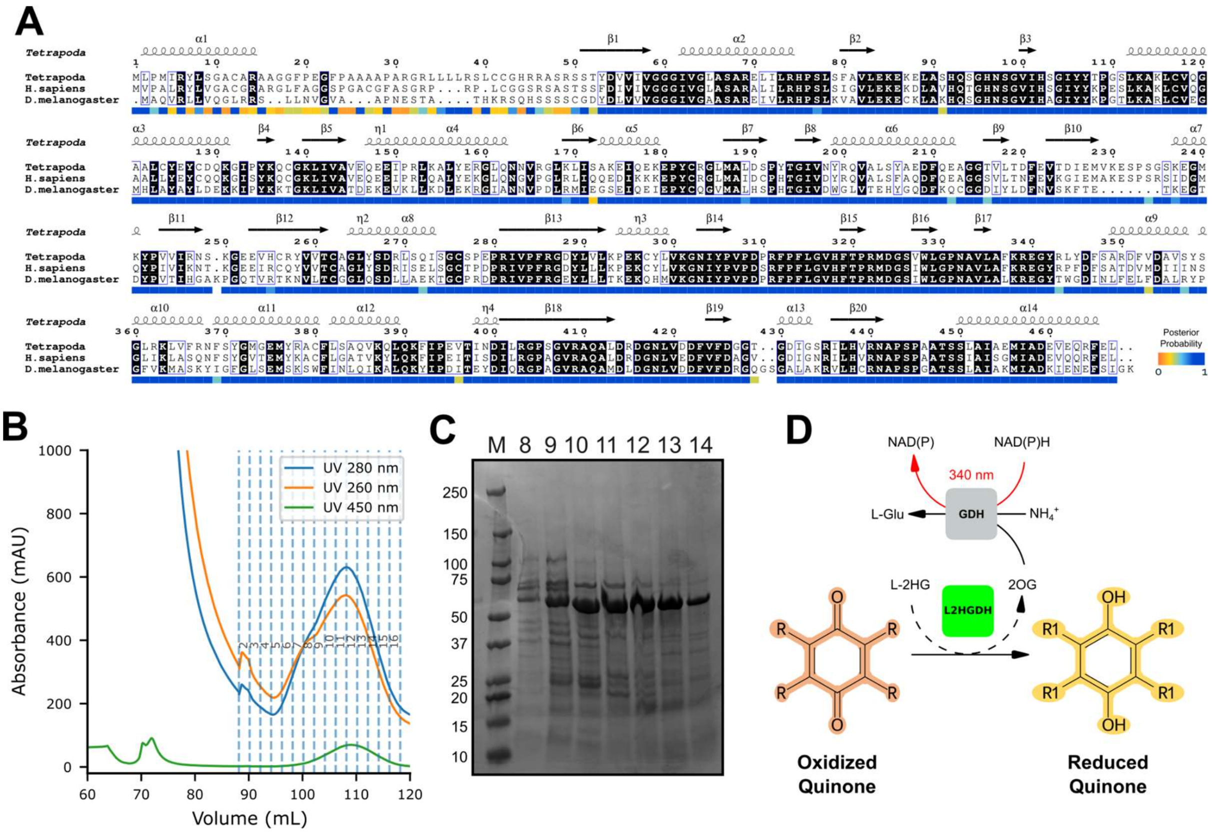

**Figure S1.** Purification of the ancestral tetrapod L2HGDH.

(**A**) Multiple sequence alignment of the reconstructed ancestral tetrapod L2HGDH with the human and *Drosophila* orthologs. Secondary structure elements are shown above the alignment based on the AlphaFold model of the ancestral enzyme. The posterior probability of the reconstructed residues is indicated in the bottom track and colored according to the blue-orange gradient shown in the scale bar. (**B-C**) Chromatogram (**B**) and SDS-PAGE (**C**) of L2HGDH purification by affinity chromatography. Fractions loaded on the gel are indicated and correspond to the chromatographic profile.

(**D**) Quinone reduction by L2HGDH was coupled to glutamate dehydrogenase (GDH) activity and monitored via NADH depletion upon addition of $NH_4Cl$. This coupled assay enables comparison across different quinone substrates under identical conditions.

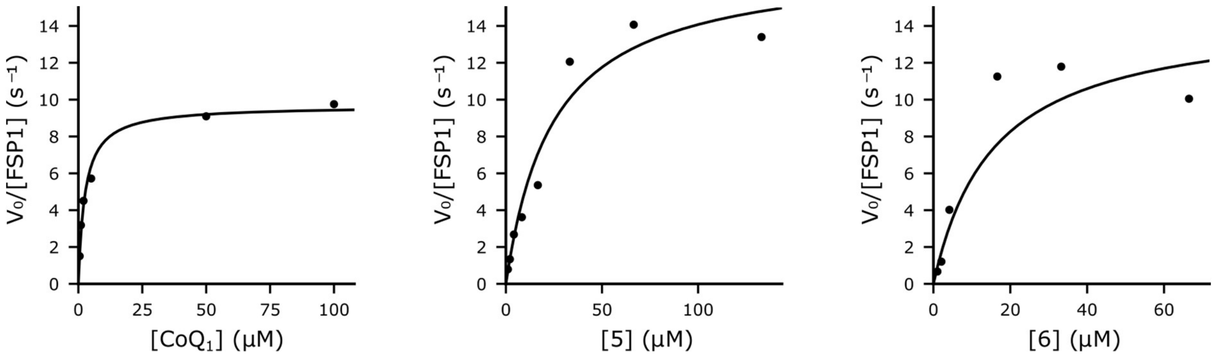

**Figure S2.** Kinetic characterization of FSP1 with quinone intermediates of the COQ metabolon.

Nonlinear Michaelis-Menten fits of FSP1 activity toward quinone substrates from the coenzyme Q biosynthetic pathway ($CoQ_1$, **6**, and **5**). Each data point represents measurements from a single independent experiment. Reactions were performed in 25 mM HEPES (pH 7.4) in the presence of NADH, the indicated quinone substrate, and FSP1.

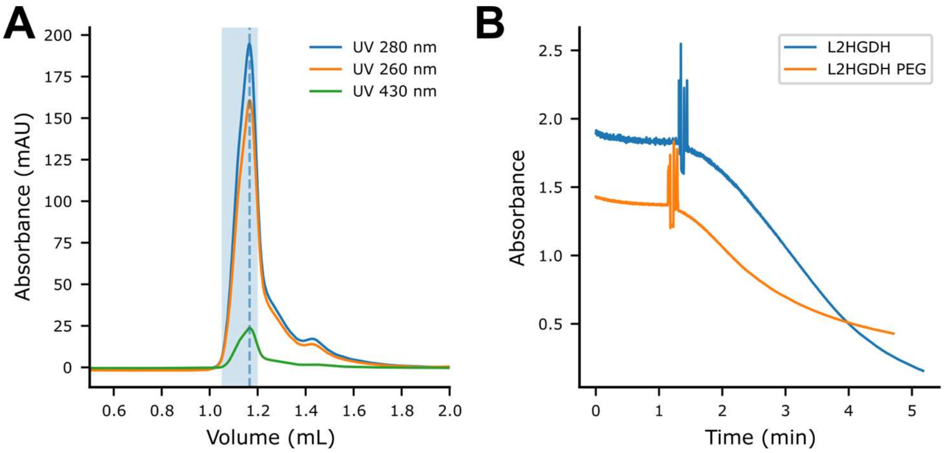

**Figure S3.** PEGylation of L2HGDH preserves quinone reductase activity.

(**A**) Size-exclusion chromatography of PEGylated L2HGDH, showing high-molecular-weight species eluting at an apparent molecular mass of approximately 900 kDa.

(**B**) Quinone reduction activity of L2HGDH and its PEGylated variant measured by monitoring NADH oxidation at 340 nm in the presence of 100 μM monoprenylated $CoQ_1$. Assays were performed using 0.1 μM enzyme. Similar reaction rates demonstrate that PEGylation preserves the catalytic activity of L2HGDH. The spike in the graph indicates the time of enzyme addition starting the reaction. Reactions were performed in 25 mM HEPES (pH 7.4) in the presence of L-2-hydroxyglutarate, $CoQ_1$, and L2HGDH.

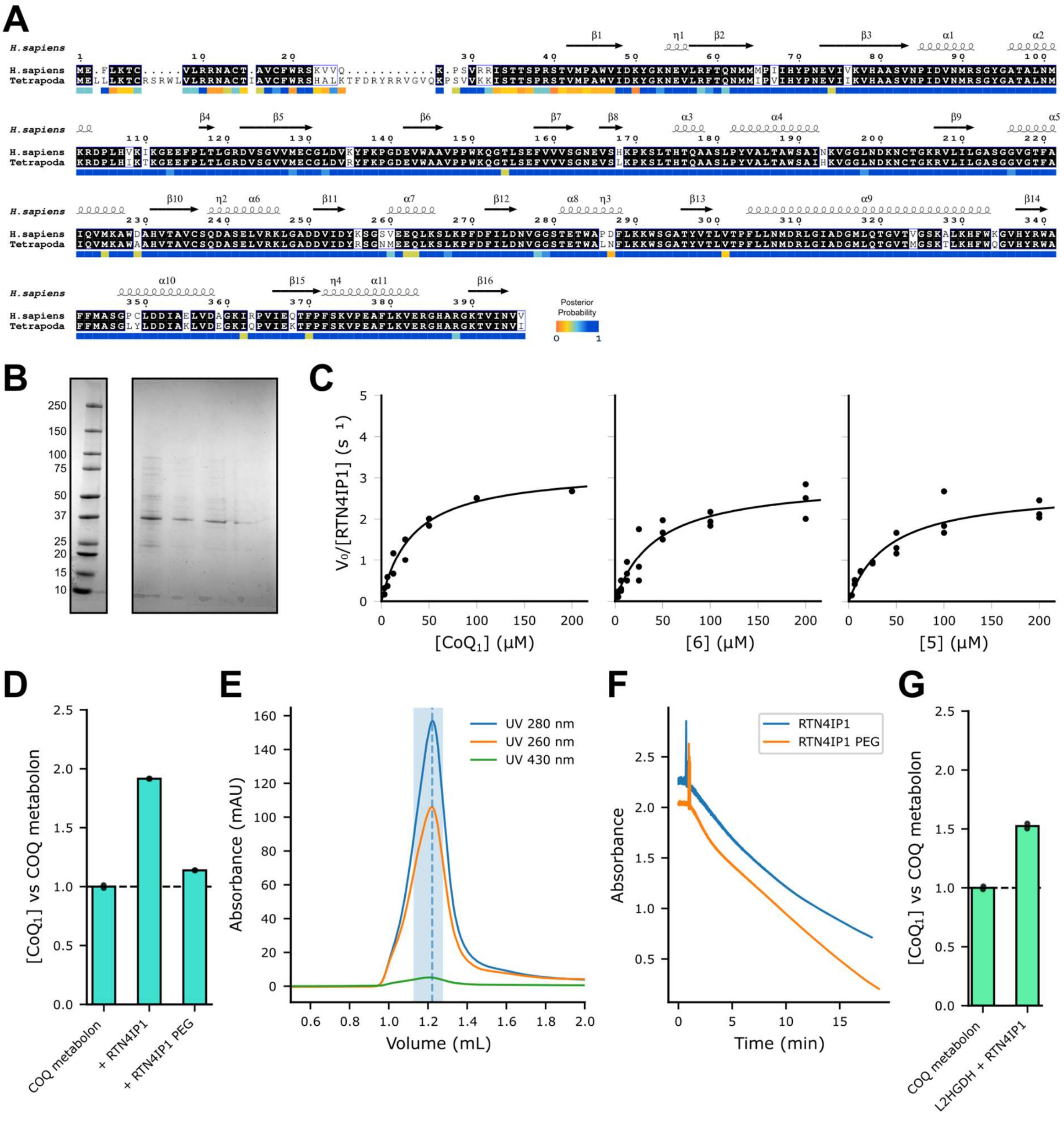

**Figure S4.** Purification and enzymatic activity of RTN4IP1

(**A**) Multiple sequence alignment of the reconstructed ancestral tetrapod RTN4IP1 with the human and ortholog. Secondary structure elements are shown above the alignment based on the crystallographic structure (PDB: 2VN8) of the human enzyme. The posterior probability of the reconstructed residues is indicated in the bottom track and colored according to the blue-orange gradient shown in the scale bar.

(**B**) SDS-PAGE of RTN4IP1 after SUMO protease cleavage purified by reversed affinity chromatography. Marker (left) and flow through fractions (right) are shown.

(**C**) Nonlinear Michaelis-Menten fits of RTN4IP1 activity toward quinone substrates from the coenzyme Q biosynthetic pathway ($CoQ_1$, **6**, and **5**). Each data point represents measurements from a single independent experiment. Reactions were performed in 25 mM HEPES (pH 7.4) in the presence of NADH, the indicated quinone substrate, and RTN4IP1.

(**D**) Bar plot showing the fold change in coenzyme $Q_1$ production from **1** in the presence of the full metabolon (control) and upon addition of RTN4IP1 or its PEGylated form (RTN4IP1 PEG).

(**E**) Size-exclusion chromatography of PEGylated RTN4IP1, showing high-molecular-weight species eluting at an apparent molecular mass of approximately 800 kDa.

(**F**) Quinone reduction activity of RTN4IP1 and its PEGylated variant measured by monitoring NADH oxidation at 340 nm in the presence of 100 µM coenzyme $Q_1$. Assays were performed using 0.1 µM enzyme. Similar reaction rates demonstrate that PEGylation preserves the catalytic activity of RTN4IP1. The spike in the graph indicates the time of enzyme addition starting the reaction. Reactions were performed in 25 mM HEPES (pH 7.4) in the presence of NADH, coenzyme $Q_1$, and RTN4IP1.
(**G**) Bar plot showing the fold change in coenzyme $Q_1$ production from **1** by the metabolon (control) upon addition of RTN4IP1 and L2HGDH.

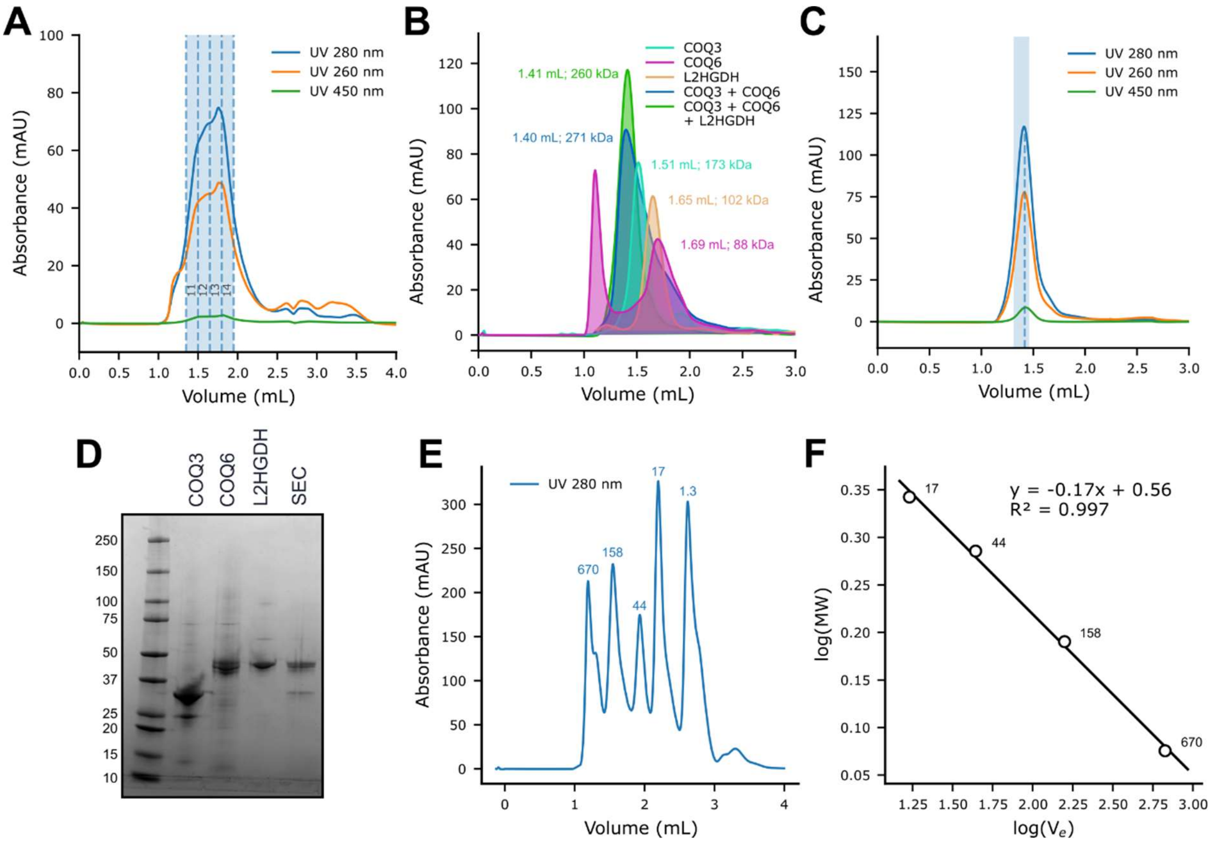

**Figure S5**. Size-exclusion chromatography analysis of interactions between COQ3, COQ6, and L2HGDH.

(**A**) Analytical size-exclusion profile of the reconstructed ancestral tetrapod COQ metabolon core, composed of COQ3, COQ4, COQ5, COQ6, COQ7 and COQ9, incubated with L2HGDH. Signal at 450 nm (flavin absorbance) reveals L2HGDH presence.

(**B**) Analytical size-exclusion chromatography of COQ3, COQ6, L2HGDH, COQ3+COQ6, and COQ3+COQ6+L2HGDH, colored according to the legend. All samples were prepared at 25 µM for each protein, and 50 µL were injected using a 100 µL loop on a Superdex 200 Increase 5/150 GL column.

(**C**) Size-exclusion chromatography of COQ3, COQ6, and L2HGDH (25 µM each) performed on a Superdex 200 Increase 5/150 GL. Absorbance was monitored at three wavelengths (280, 260, and 450 nm). The fraction used for cryoEM is highlighted in light blue. The peak is highlighted with blue dashed line.

(**D**) SDS-PAGE of individual proteins (COQ3, COQ6, L2HGDH) and of the SEC fraction indicated in panel **B**, confirming co-elution of all three components.

(**E**) Size-exclusion chromatography of molecular-weight standards used to estimate the apparent molecular weight of protein complexes from their elution volume. Standards were, in elution order, thyroglobulin from bovine thyroid (670 kDa), γ-globulin from bovine blood (158 kDa), ovalbumin from chicken egg white (44 kDa), myoglobin from horse skeletal muscle (17 kDa), and vitamin B12 (1.35 kDa).

(**F**) Calibration curve generated from the elution volumes of the standards shown in panel **D**, used to estimate the apparent molecular weights of the complexes shown in panel **A**.

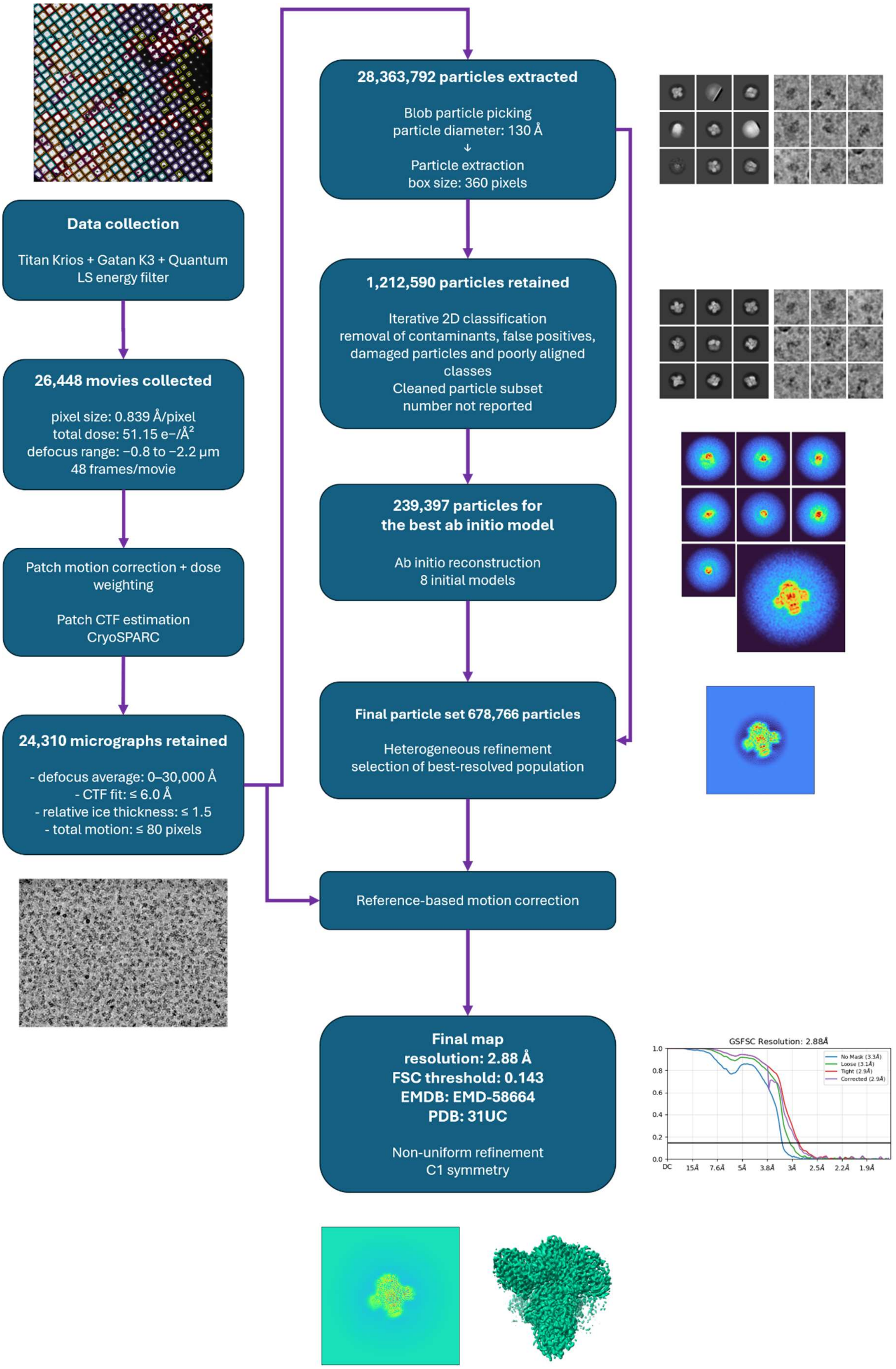
Data collection
Titan Krios + Gatan K3 + Quantum LS energy filter
26,448 movies collected
pixel size: 0.839 Å/pixel
total dose: 51.15 e−/Å²
defocus range: −0.8 to −2.2 µm
48 frames/movie
Patch motion correction + dose weighting
Patch CTF estimation
CryoSPARC
24,310 micrographs retained
- defocus average: 0–30,000 Å
- CTF fit: ≤ 6.0 Å
- relative ice thickness: ≤ 1.5
- total motion: ≤ 80 pixels
28,363,792 particles extracted
Blob particle picking
particle diameter: 130 Å
↓
Particle extraction
box size: 360 pixels
1,212,590 particles retained
Iterative 2D classification
removal of contaminants, false positives, damaged particles and poorly aligned classes
Cleaned particle subset
number not reported
239,397 particles for the best ab initio model
Ab initio reconstruction
8 initial models
Final particle set 678,766 particles
Heterogeneous refinement
selection of best-resolved population
Reference-based motion correction
Final map
resolution: 2.88 Å
FSC threshold: 0.143
EMDB: EMD-58664
PDB: 31UC
Non-uniform refinement
C1 symmetry
GSFSC Resolution: 2.88Å
No Mask (3.3Å)
Loose (3.1Å)
Tight (2.9Å)
Corrected (2.9Å)
1.0
0.8
0.6
0.4
0.2
0.0
DC
15Å
7.6Å
5Å
3.8Å
3Å
2.5Å
2.2Å
1.9Å

**Figure S6**. Pipeline of CryoEM data collection and processing.

Scheme of the cryoSPARC workflow used to obtain the final deposited map (EMDB: EMD-58664) from the data collection at CM01 (ESRF, Grenoble) [58]

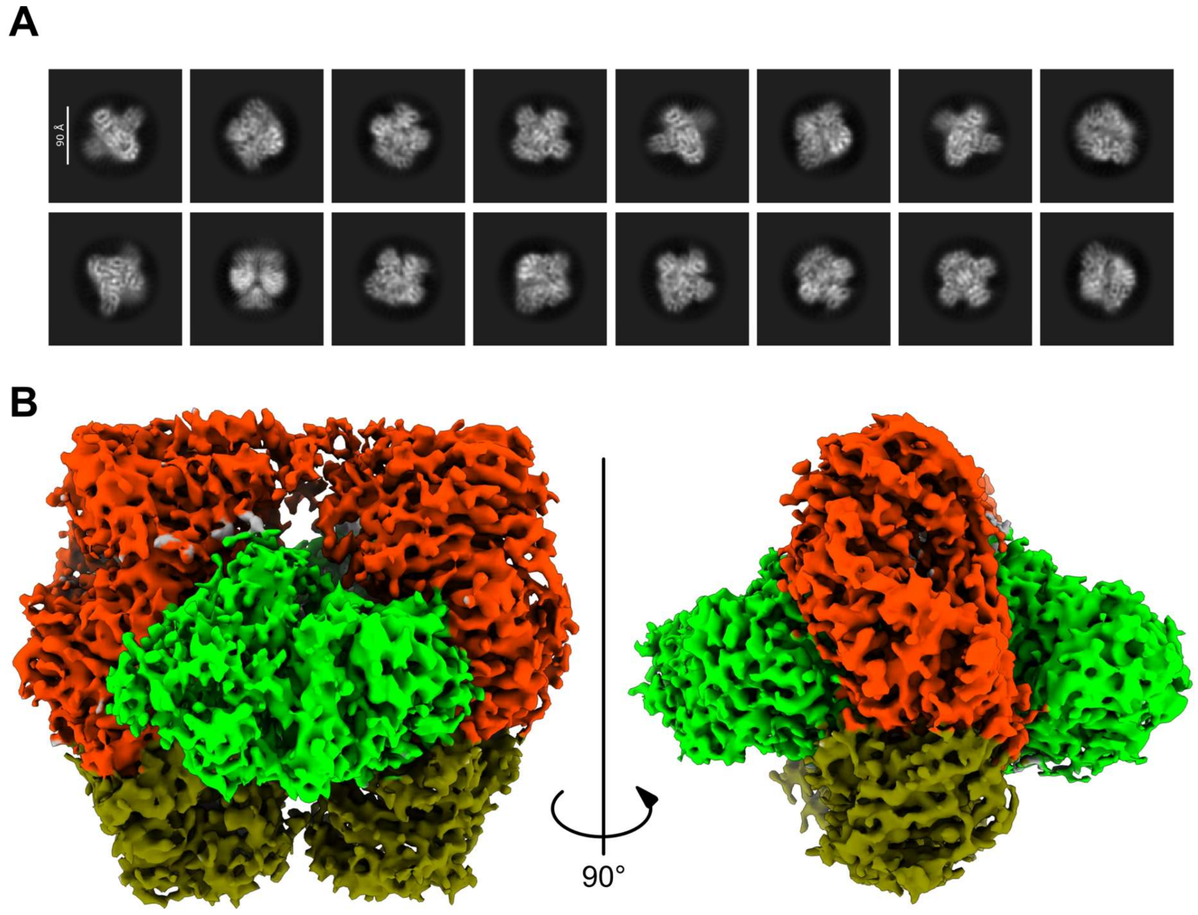

**Figure S7.** Cryo-EM reconstruction of the COQ3-COQ6-L2HGDH assembly.

(**A**) Representative cleaned 2D class averages of the complex.

(**B**) CryoEM reconstruction of the COQ3-COQ6-L2HGDH complex, revealing a hexameric assembly. Subunits are colored as in **Figure 1**. The map is shown at a contour level of 2 σ.

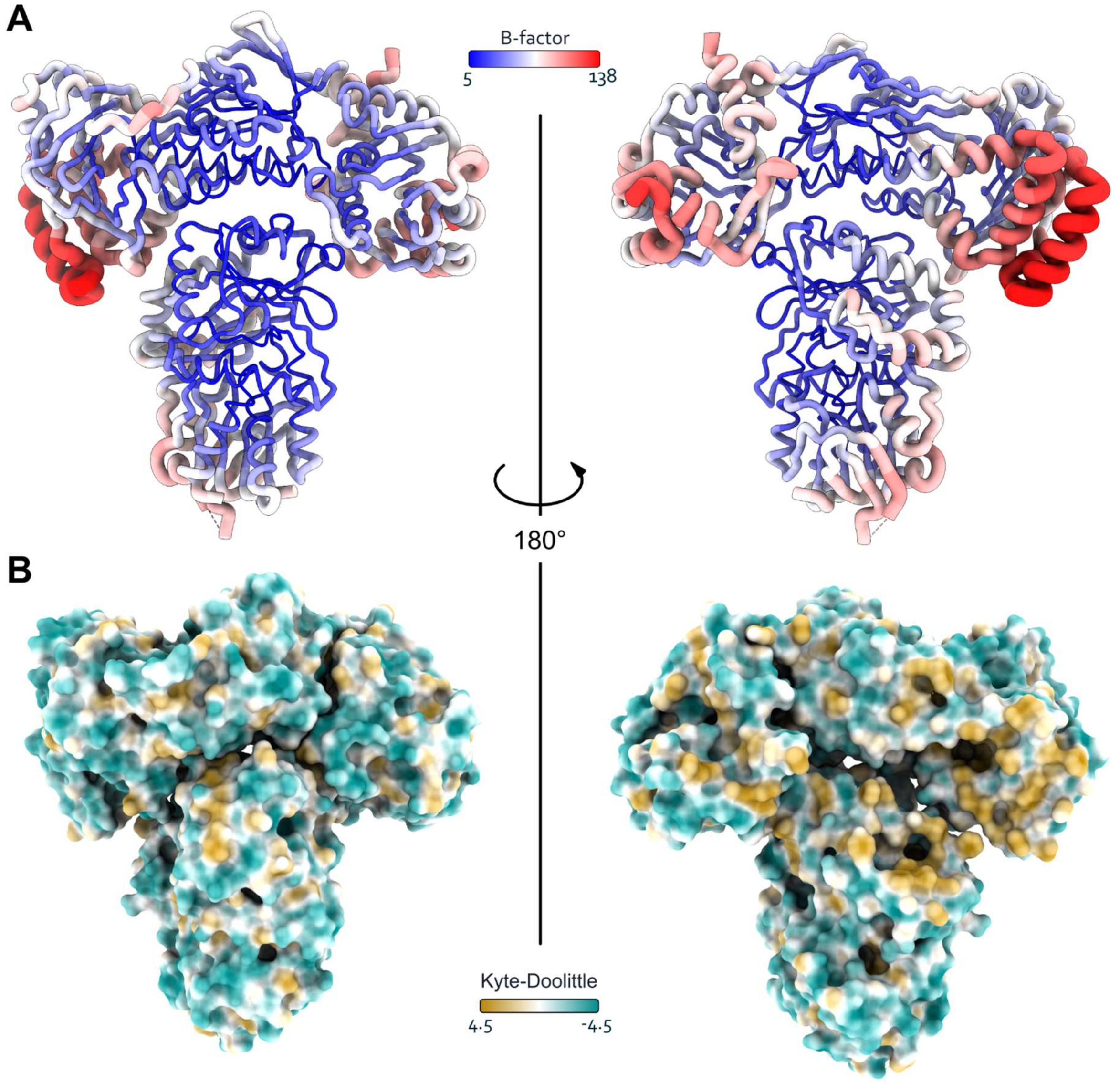

**Figure S8.** B-factor distribution of the COQ3-COQ6-L2HGDH trimer.

(**A**) Cartoon tube representation of the trimer colored according to B-factor values using the blue-white-red scale shown. Regions with higher B-factors are additionally highlighted by increased tube thickness. The highest B-factor values are observed in the regions contacting the second, less well-resolved trimer in the hexameric reconstruction.

(**B**) surface representation of the same orientation of panel **A** showing colored by hydrophobicity scale (Kyte-Doolittle)

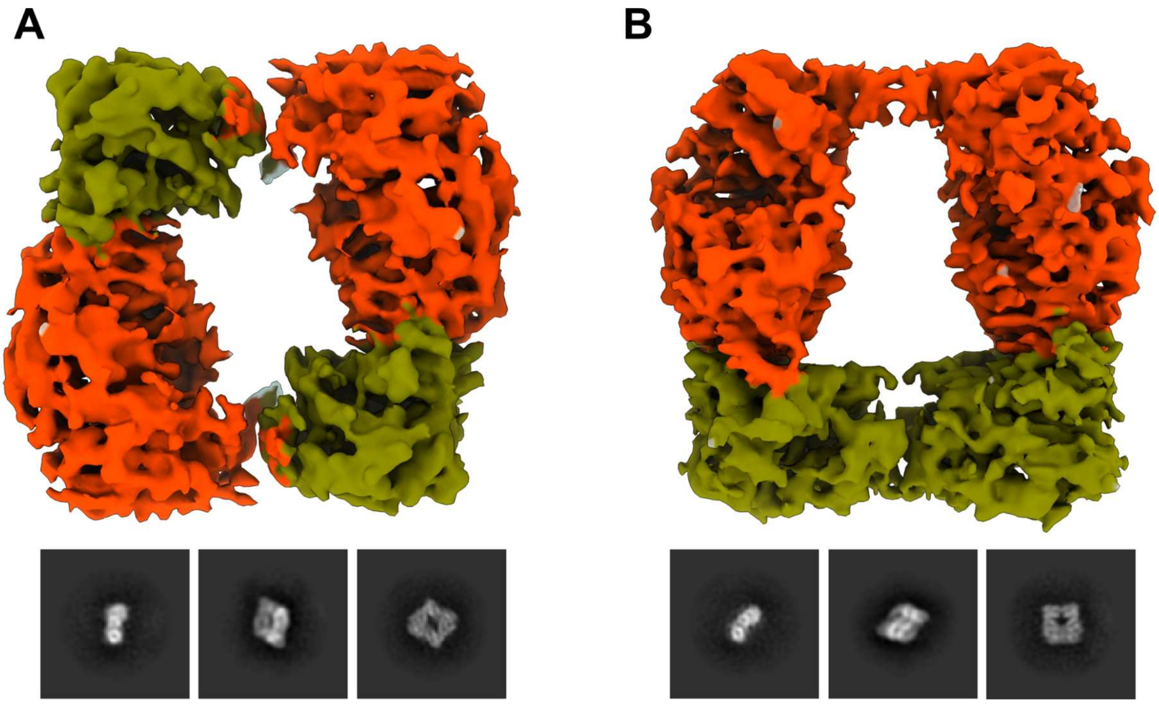

**Figure S9**. Cryo-EM reconstruction of the COQ3-COQ6 assemblies.

(**A-B**) Cryo-EM models (top) and 2D class averages (bottom) of the COQ3-COQ6 complex in the absence of L2HGDH reveal a stable heterodimer that assembles into a dimer of dimers adopting two alternative conformations: head-to-tail (**A**) and head-to-head (**B**).

A

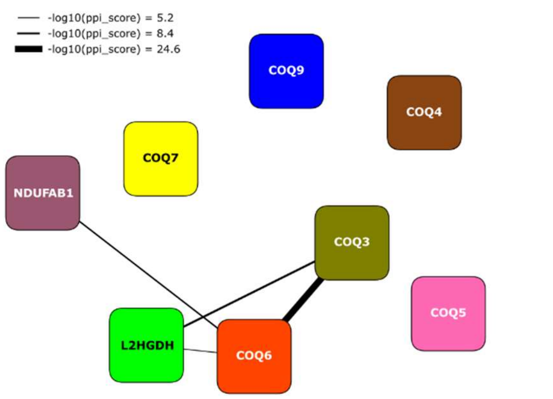

B

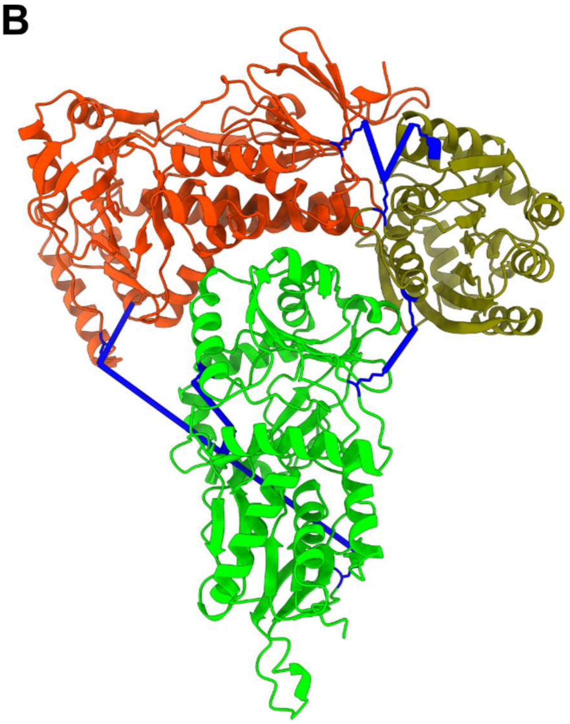

C

| Residue A | Residue B | Distance (Å) |
|---|---|---|
| COQ3 LYS 196 NZ | COQ3 LYS 143 NZ | 10.155 |
| COQ3 LYS 143 NZ | COQ6 LYS 64 NZ | 10.531 |
| L2HGDH LYS 155 NZ | L2HGDH LYS 109 NZ | 10.612 |
| COQ3 LYS 134 NZ | L2HGDH LYS 132 NZ | 12.839 |
| COQ6 LYS 325 NZ | COQ6 LYS 459 NZ | 13.571 |
| L2HGDH LYS 246 NZ | COQ6 LYS 325 NZ | 57.329 |

**Figure S10.** Mitochondrial crosslinking of COQ metabolon components and external interactors.

(**A**) Network of intra- and inter-metabolon interactions involving COQ metabolon components identified in published cross-link assisted spatial proteomics in the mitochondrial membrane[35]. Edge thickness reflects the protein-protein interaction (PPI) score, as indicated in the figure legend.
(**B**) Cartoon representation of Boltz-2 models of human COQ3, COQ6, and L2HGDH superimposed to the tetrapod ternary complex. Lysine residues involved in crosslinking are highlighted as blue sticks and corresponding crosslinks are shown as blue lines[35].
(**C**) Table of crosslinked residues shown in panel **B**, including the corresponding Nε-Nε distances in angstroms. Distances compatible with the DSBSO crosslinker are highlighted in light blue.

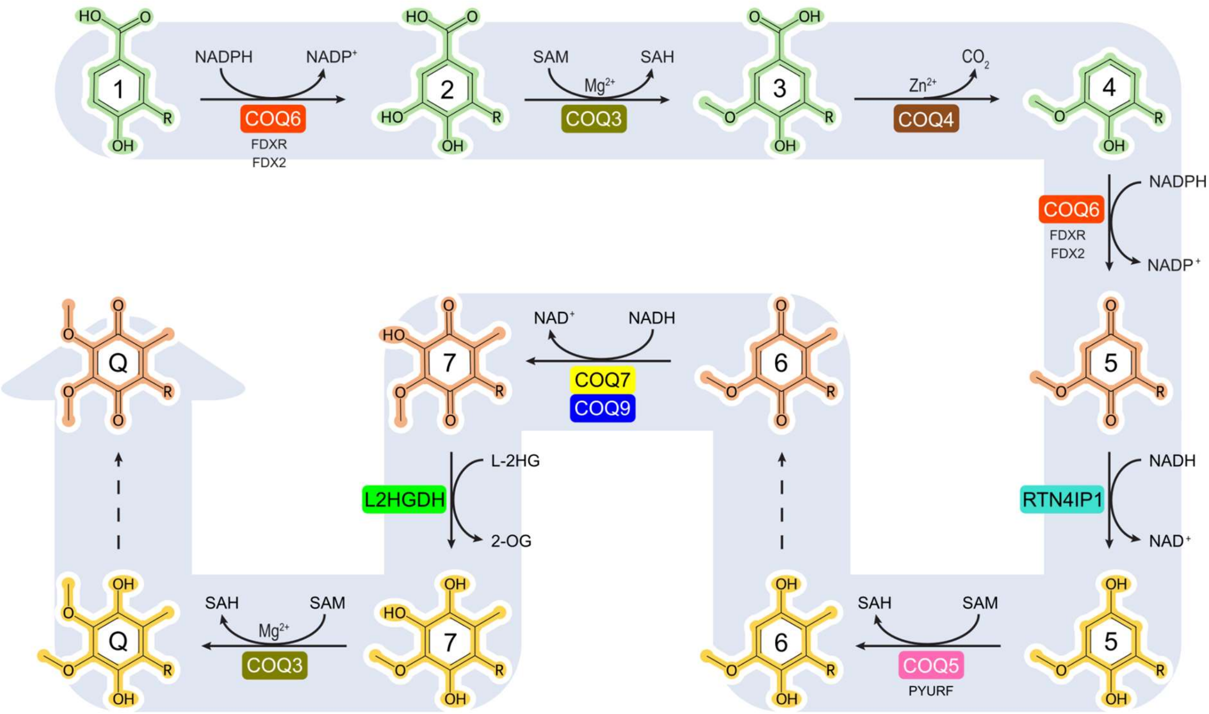

**Figure S11**. Updated model of coenzyme Q biosynthesis within the COQ metabolon.

Schematic representation of the coenzyme Q biosynthetic pathway and the associated enzymes. Intermediates are numbered from **1** to **7**, and the final product coenzyme Q is abbreviated as Q. Compounds are colored to distinguish the early non-quinone phase (green) from the late quinone phase, with oxidized intermediates shown in salmon and reduced intermediates in yellow. RTN4IP1 and L2HGDH, act upstream of the two methyltransferase reactions catalyzed by COQ5 and COQ3, respectively, within the quinone phase. Dashed lines indicate spontaneous reoxidation of intermediates or the final product.

| Enzyme | Substrate | $k_{cat}$ ($s^{-1}$) | $K_M$ (μM) | $k_{cat}$ / $K_M$ ($M^{-1}$ $s^{-1}$) |
|---|---|---|---|---|
| **L2HGDH** | **5** | 8.45 ± 0.23 | 12.68 ± 1.09 | 6.66 x 1e5 ± 0.75 x 1e5 |
| | **6** | 11.19 ± 0.36 | 14.60 ±1.41 | 7.66 x 1e5 ± 0.99 x 1e5 |
| | **$CoQ_1$** | 18.95 ± 0.99 | 26.223 ± 3.68 | 7.23 x 1e5 ± 1.39 x 1e5 |
| **FSP1** | **5** | 17.54 ±1.17 | 24.59 ± 4.62 | 7.13x1e5 ± 1.82 x 1e5 |
| | **6** | 13.40 ± 1.25 | 8.05 ± 2.6 | 16.65x1e5 ± 6.95 x 1e5 |
| | **$CoQ_1$** | 9.67 ± 0.21 | 2.56 ± 0.21 | 37.8x1e5 ± 3.98 x 1e5 |
| **RTN4IP1** | **5** | 2.73 ± 0.15 | 41.56 ± 16.41 | 0.66x1e5 ± 0.75 x 1e5 |
| | **6** | 2.95 ± 0.16 | 42.24 ±6.18 | 0.70x1e5 ± 0.14 x 1e5 |
| | **$CoQ_1$** | 3.21 ± 0.09 | 32.36 ± 2.42 | 0.99x1e5 ± 0.10 x 1e5 |

**Table S1.** Kinetic parameters of L2HGDH, FSP1 and RTN4IP1 with quinone substrates.

Kinetic parameters ($k_{cat}$, $K_M$, and catalytic efficiency, $k_{cat}/K_M$) measured with $CoQ_1$ and quinone intermediates of the coenzyme Q biosynthetic pathway (**6** and **5**). Values represent mean ± standard error from nonlinear Michaelis-Menten fitting.

| | PDB: 31UC<br>EMDB: EMD-58664 | EMDB: EMD-58668<br>Head-to-tail | EMDB: EMD-58667<br>Head-to-head |
|---|---|---|---|
| **Data collection and processing** | COQ3-COQ6-L2HGDH | COQ3-COQ6 | |
| Microscope | Titan Krios | | |
| Magnification | 105,000 | 81,000 | |
| Voltage (kV) | 300 | | |
| Detector | Gatan K3 | | |
| Electron exposure ($e^-/Å^2$) | 51.15 | 50.55 | |
| Defocus range (um) | from -0.8 to 2.2 | from -1.0 to 2.2 | |
| Pixel size (Å) | 0,839 | 1.06 | |
| Movies (no.) | 26,448 | 17,828 | |
| Symmetry imposed | C1 | C2 | |
| Final particle images (no.) | 678,766 | 123,481 | 135,927 |
| Map resolution (Å) | 2.88 | 3.95 | 3.67 |
| FSC threshold | 0.143 | | |
| **Refinement** | | | |
| Initial model used | COQ3 crystal structure and Boltz2 predictions | | |
| **Model composition (#)** | | | |
| Non-hydrogen atoms | 8,227 | | |
| Protein residues | 1,051 | | |
| Ligands | FAD:1 | | |
| **B factor ($Å^2$)** | | | |
| Protein (min/max/mean) | 0.00/131.43/32.97 | | |
| Ligand (min/max/mean) | 0.42/41.11/13.26 | | |
| **Bonds (RMSD)** | | | |
| Length (Å) (# > 4 σ) | 0.003 | | |
| Angles (Å) (# > 4 σ) | 0.553 | | |
| CC_mask | 0.80 | | |
| **Validation** | | | |
| **Ramachandran plot** | | | |
| Residues favored (%) | 96.06 | | |
| Residues disallowed (%) | 0.00 | | |

| | |
|---|---|
| MolProbity score | 2.03 |
| Clashscore | 5.63 |
| Rotamers outliers (%) | 3.74 |

**Table S2.** CryoEM density and model processing and validation parameters

| PDB | 31QD |
|---|---|
| Space group | $C222_1$ |
| Unit cell axes (Å) | 90.5<br>147.4<br>160.3 |
| Resolution (Å) | 3.18 |
| $R_{sym}$ [a, b] (%) | 23.7 (182.5) |
| $CC_{1/2}$ [b] (%) | 99.5 (53.7) |
| Completeness [b] (%) | 99.9 (99.9) |
| Unique reflections | 18408 |
| Redundancy [b] | 10.6 (10.8) |
| I/σ [b] | 8.4 (1.5) |
| N. of non-hydrogen atoms<br>protein<br>ligands<br>water | <br>5401<br>172<br>27 |
| Average B factor for protein/ligands | 84.93/114.34 |
| $R_{cryst}$ [c] (%) | 19.47 |
| $R_{free}$ [c] (%) | 27.56 |
| Rms bond length (Å) | 0.0059 |
| Rms bond angles (Å) | 1.5630 |

**Table S3.** Data collection and refinement statistics for the crystal structure of tetrapod ancestral COQ3 in complex with intermediate **2**, $Mg^{2+}$ and SAH.

[a] $Rsym=\sum|I_i-\langle I\rangle|/\sum I_i$, where $I_i$ is the intensity of $i^{th}$ observation and $\langle I\rangle$ is the mean intensity of the reflection.

[b] Values in parentheses are for reflections in the highest resolution shell.

[c] $R_{cryst}=\sum|F_{obs}-F_{calc}|/\sum|F_{obs}|$ where $F_{obs}$ and $F_{calc}$ are the observed and calculated structure factor amplitudes, respectively. $R_{cryst}$ and $R_{free}$ were calculated using the working and test sets, respectively.
